# Phosphoproteome profiling uncovers a key role for CDKs in TNF signaling

**DOI:** 10.1101/2020.11.04.368159

**Authors:** Maria C Tanzer, Isabell Bludau, Che A Stafford, Veit Hornung, Matthias Mann

## Abstract

Tumor necrosis factor (TNF) is one of the few cytokines successfully targeted by therapies against inflammatory diseases. However, blocking this well studied and pleiotropic ligand can cause dramatic side-effects. We reasoned that a systems-level proteomic analysis of TNF signaling could dissect its diverse functions and offer a base for developing more targeted therapies. Combining phosphoproteomics time course experiments with subcellular localization and kinase inhibitor analysis identifies functional modules of phosphorylations. The majority of regulated phosphorylations could be assigned to an upstream kinase by inhibiting master kinases and spatial proteomics revealed phosphorylation-dependent translocations of hundreds of proteins upon TNF stimulation. Phosphoproteome analysis of TNF-induced apoptosis and necroptosis uncovered a key role for transcriptional cyclin-dependent kinase (CDK) activity to promote cytokine production and prevent excessive cell death downstream of the TNF signaling receptor. Our comprehensive interrogation of TNF induced pathways and sites can be explored at http://tnfviewer.biochem.mpg.de/.

**Highlights:**

- Distinct phosphorylation events mark early and late TNF signaling
- Inhibition of master kinases reveals TNF stimulation dependent kinase-substrate relations
- TNF induces phosphorylation-dependent spatial rearrangement of hundreds of proteins
- CDK kinase activity promotes TNF-induced cytokine expression and inhibits cell death
- CDK12/13 inhibitors have potential as anti-inflammatory agents

## Introduction

Post-translational modifications (PTMs) such as phosphorylation govern the activation, strength, and timing of immune signaling pathways. The interplay between kinases and phosphatases results in the rapid addition and removal of phosphates, providing exquisitely precise control of signaling events. In addition to a small number of key phosphorylation sites with switch-like functions, there is a vast number of phosphorylation events that fine-tune cellular responses (Taniguchi et al., 2006) (Ochoa et al., 2020).

The TNF pathway is a pivotal proinflammatory signaling cascade that relies heavily on protein phosphorylation, the extent of which has been revealed by phosphoproteomic studies (Wagner et al., 2016, Mohideen et al., 2017, Zhong et al., 2014, Krishnan et al., 2015, Welz et al., 2019, Cantin et al., 2006). Much is already known about key mechanistic events and we briefly recapitulate those that are pertinent to our study. TNF binds to its receptor (TNFR1), which leads to the recruitment of the adaptor proteins TNF receptor-associated factor 2 (TRAF2), Tumor necrosis factor receptor type 1-associated death domain protein (TRADD), and the Receptor-interacting serine/threonine-protein kinase 1 (RIPK1), which play critical roles in the decision between cell death, survival, and inflammation (Hsu et al., 1996, Ting et al., 1996). This complex is then ubiquitylated, allowing recruitment and activation of the master kinase TGF-β activated kinase-1 (TAK1), which then activates MAPK- and NF-κB signaling by phosphorylating several MAPK kinases and IκB kinases (IKK1/2) (Wertz and Dixit, 2010, Silke, 2011). Both pathways are crucial for the upregulation of many target genes, including a range of cytokines and pro-survival factors (Hayden and Ghosh, 2008). One such pro-survival factor is the cellular FLICE-like inhibitory protein (cFLIP), which is required to inhibit caspase-8 activity and prevent cell death (Budd et al., 2006). Disruption of these phosphorylation-driven signaling cascades strongly perturbs gene activation, leading to cell death and inflammation (Mihaly et al., 2014). Besides their roles in transcriptional activation, IKK2 and MAPK-activated protein kinase 2 (MK2) directly phosphorylate RIPK1, inhibiting cell death by altering RIPK1’s kinase and adaptor function (Dondelinger et al., 2015, Jaco et al., 2017). TNF-induced cell death can either be caspase-dependent (apoptosis) or independent (necroptosis). Necroptosis induction occurs in the absence or upon inhibition of caspase-8. It involves RIPK1 autophosphorylation and activation of RIPK3, which subsequently phosphorylates and activates the Mixed lineage kinase domain like pseudokinase (MLKL) leading to plasma membrane permeabilization and cell death (Sun et al., 2012, Hildebrand et al., 2014, Murphy et al., 2013). Necroptosis is considered to be inflammatory (Kaczmarek et al., 2013). In line with this concept, we previously showed that MLKL activation leads to the release of proteases, and other intracellular proteins (Tanzer et al., 2020).

Clinically targeting TNF is highly successful in treating inflammatory pathologies (Croft et al., 2013, Bradley, 2008). Most anti-inflammatory therapies, including TNF-blocking antibodies, act by inhibiting cytokines that drive inflammation, leading to a complete abrogation of all downstream signaling events. Considering the pleiotropic functions of cytokines – especially in fighting infection and regulating a range of important signaling pathways – it is not surprising that anti-inflammatory therapies often cause severe side effects (Bradley, 2008). In light of this, targeting specific phosphorylation events by inhibiting certain kinases or phosphatases could allow more precise manipulation of disease pathways while retaining signal transduction required for homeostasis. However, such an approach requires an in-depth knowledge of kinases, phosphatases, substrates, and their signaling dynamics, which is currently unavailable. Furthermore, the complexity and interplay of these events further make analysis by classical cell biology tools difficult. We therefore reasoned that systems-level approaches like proteomics and phosphoproteomics provide the required global cellular perspective on phosphorylation dynamics.

To investigate the TNF-regulated phosphoproteome we combined TNF-biology in myeloid cell systems with a high sensitivity phospho-enrichment protocol (Humphrey et al., 2018, Humphrey et al., 2015) and state of the art mass spectrometry. We specifically made use of the data completeness of the data-independent acquisition mode (Ludwig et al., 2018). We employed time course and spatial proteomics to elucidate signaling events downstream of the TNF receptor. Based on this in-depth data, we functionally probed the impact of master kinases and TNF-induced cell death on the global TNF-phosphoproteome, revealing a plethora of signaling events not previously connected to TNF. Our findings offer a comprehensive resource of phosphorylation events regulated upon TNF-stimulation and TNF-induced cell death, which is available to the community at http://tnfviewer.biochem.mpg.de/. We provide evidence for TNF-mediated cross-talk with other innate immune signaling pathways and identify a role for CDK kinase activity in TNF-induced cell death.

## Results

### Temporal analysis of TNF-induced phosphorylation events enables separation of early and late signaling events

To acquire a systems-based view of phosphorylation events and their kinetics downstream of the TNF receptor complex, we treated the myeloid cell line U937 with TNF over a time course of fifteen s to one hour (**Figure 1A, B**). We analyzed phosphopeptides in a data-independent acquisition (DIA) mode and detected 28,000 class I phosphosites (localization probability > 75 %) (**Figure 1C**). Overall, significantly changing TNF-induced phosphosites peaked at 15 min, but we detected significant upregulations already at 90 s after treatment (**Figure 1D, E**). The large majority returned to baseline by 60 min (**Figure 1E, F**). Many of these transiently modified proteins are involved in NF-κB- and pattern recognition signaling. In contrast, a small cluster of phosphosites remain upregulated even at 60 min post TNF stimulation and were located on proteins involved in transcription (**Figure 1F**). We categorized phosphosites into early (significantly regulated up to 5 min of stimulation), middle (at 15 min), and late (at 60 min) events (**Figure 1G, H**). Fisher’s exact test on the different temporal sections revealed the dynamic regulation of the respective cellular processes. Within five min of TNF-stimulation, the phosphorylation status of proteins involved in regulation of vesicle fusion and myeloid cell differentiation was increased, while terms involving the RIG-I-, NF-κB -, TRIF- and MYD88 pathways, as well as GTPase activation, followed at 15 min (**Figure 1G, H**). Terms related to transcription were regulated throughout the time course - most strongly at the latest time point, consistent with transcription being the most downstream process of the TNF signaling cascade. Motif enrichment analysis revealed a dynamic activation of different kinases along the time course (see Methods, **Figure 1I**). CDK1/2 and PLK1/3 motifs were downregulated, whereas IKBKB (IKK2) and PRKAA1/2 motifs were upregulated at early time points. Phosphorylations on one protein could present with different kinetics. For example, of the quantified phosphosites of INPP5D, a phosphatase involved in immune-signaling, S886 and T1108 peaked at 8 min whereas T963 and T971 only peaked at 60 min, which suggests that different regulators act on one protein (**Supplementary Figure 1A**).

**Figure 1:**
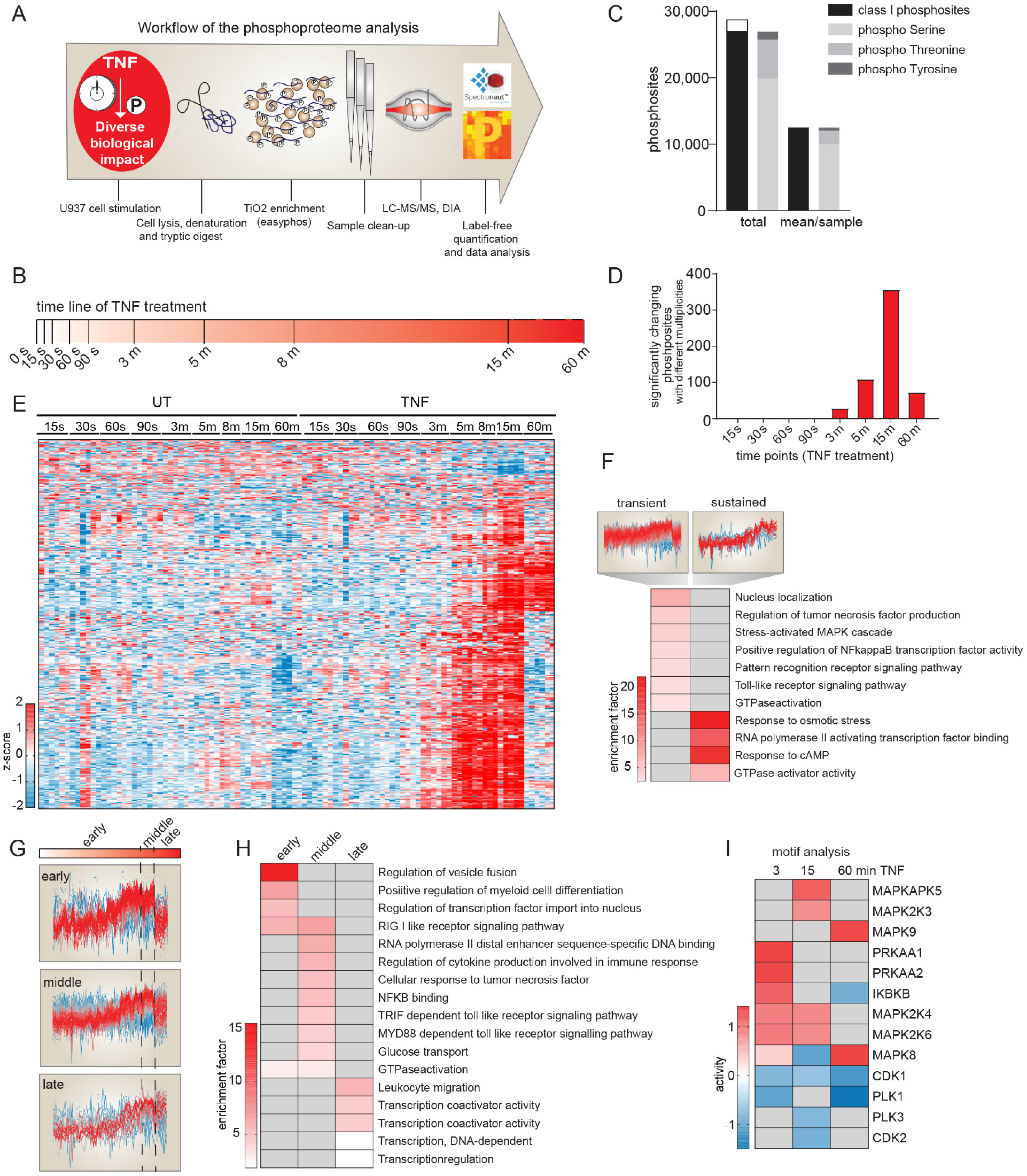
Temporal dissection of TNF-regulated phosphorylation events. (**A**) Workflow of the phosphoproteome analysis. (**B**) Schematic overview of the time points used for TNF treatment (100 ng/ml). (**C**) Bar graph demonstrating number of detected phosphosites in U937 cells. Class I sites (in black) are phosphosites with a localization probability of > 75%. (**D**) Histogram depicting the numbers of significantly changing phosphosites along 8 different time points compared to the respective untreated control (FDR < 0.05; see STAR Method for 8 min time point). (**E**) Heatmap of significantly changing phosphosites along the time course compared to their respective untreated controls (Student’s t-test; FDR < 0.05). Z-scores of various time points and the 4-6 replicates in red for intensities higher than the mean and in blue for intensities lower than the mean of respective phosphosites across all samples. (**F**) Fisher’s exact test of transient phosphosites induced upon TNF, which are downregulated after 15 min and sustained phosphosites, which are still upregulated at 60 min of treatment (p < 0.002). The red scale resembles enrichment, while grey represents missing values/no enrichment. The profiles are color coded according to their distance from the respective cluster center (red is close to center, blue is further away from center). (**G**) Z-scored profiles of early (significantly upregulated up to 5 min of TNF treatment compared to the respective untreated control (Student’s t-test FDR < 0.05)), middle (significantly upregulated only at 15 min of TNF stimulation), and late phosphorylations (significantly upregulated only at 60 min of TNF treatment). Each profile is color coded according to its distance from the respective cluster center (red is close to center, blue is further away from center). (**H**) Fisher’s exact test of early, middle, and late regulated phosphosites (p < 0.01). The enrichment factor increases along the red gradient. Grey means not regulated. **I**) Motif analysis (see Methods) reveals kinase activity regulation at three different time points of TNF stimulation. Red scale represents increased kinase activity, while blue represents decreased activity and grey means not regulated.

With the time course on U937 cells in hand, we selected the 15 min time point to interrogate TNF signaling in another relevant cell system, murine bone marrow derived macrophages (BMDMs) (**Supplementary Figure 1C, D**). While we observed many classical TNF signaling phosphorylations, surprisingly, many proteins involved in other immune response pathways like MAVS (S222, S419) in U937 cells or NLRC4 (S533) and OAS3 (S384) in BMDMs were also dynamically phosphorylated (**Supplementary Figure 1B, C, D; Supplementary Table 1**). Terms like peptidoglycan response, TLR signaling, and response to exogenous dsRNA were significantly enriched, suggesting a cross-priming function for TNF upon infection (Fisher’s exact test **Supplementary Figure 1E, F**).

### Kinase hub inhibition unravels kinase-substrate relations upon TNF stimulation

While our analysis of phosphorylation dynamics enabled the temporal dissection of phosphorylation events, identification of specific kinase-substrate relationships remained challenging due to the simultaneous activation of many downstream kinases. To address this, we targeted key kinases such as TAK1 using specific inhibitors (**Supplementary Figure 2B**). TAK1 plays a crucial, upstream role in activating NF-κB signaling by phosphorylating IKKs (**Figure 2A**). This pathway is required for the upregulation of pro-survival target genes and is thereby essential for cell survival upon TNF stimulation (**Figure 2B**). TAK1 is also important for the activation of the MAPK pathway by phosphorylating MAPKs like p38, MEK, and JNK (**Figure 2A**). While TNF stimulation alone induced the regulation of hundreds of phosphorylation events, inhibiting TAK1 almost completely abrogated its effect (**Figure 2C; Supplementary Figure 2A**). This confirms the upstream role of TAK1 in TNF signaling using a global phospho-proteomics readout. In contrast, inhibition of downstream kinases such as IKK1/2, p38, MEK1/2, and JNK had a modest to intermediate effect on TNF-regulated phosphosites (**Figure 2C; Supplementary Figure 2A**). Inhibiting p38 had the strongest impact, reducing 76% of the TNF-upregulated phosphosites (**Supplementary Figure 2A**).

**Figure 2:**
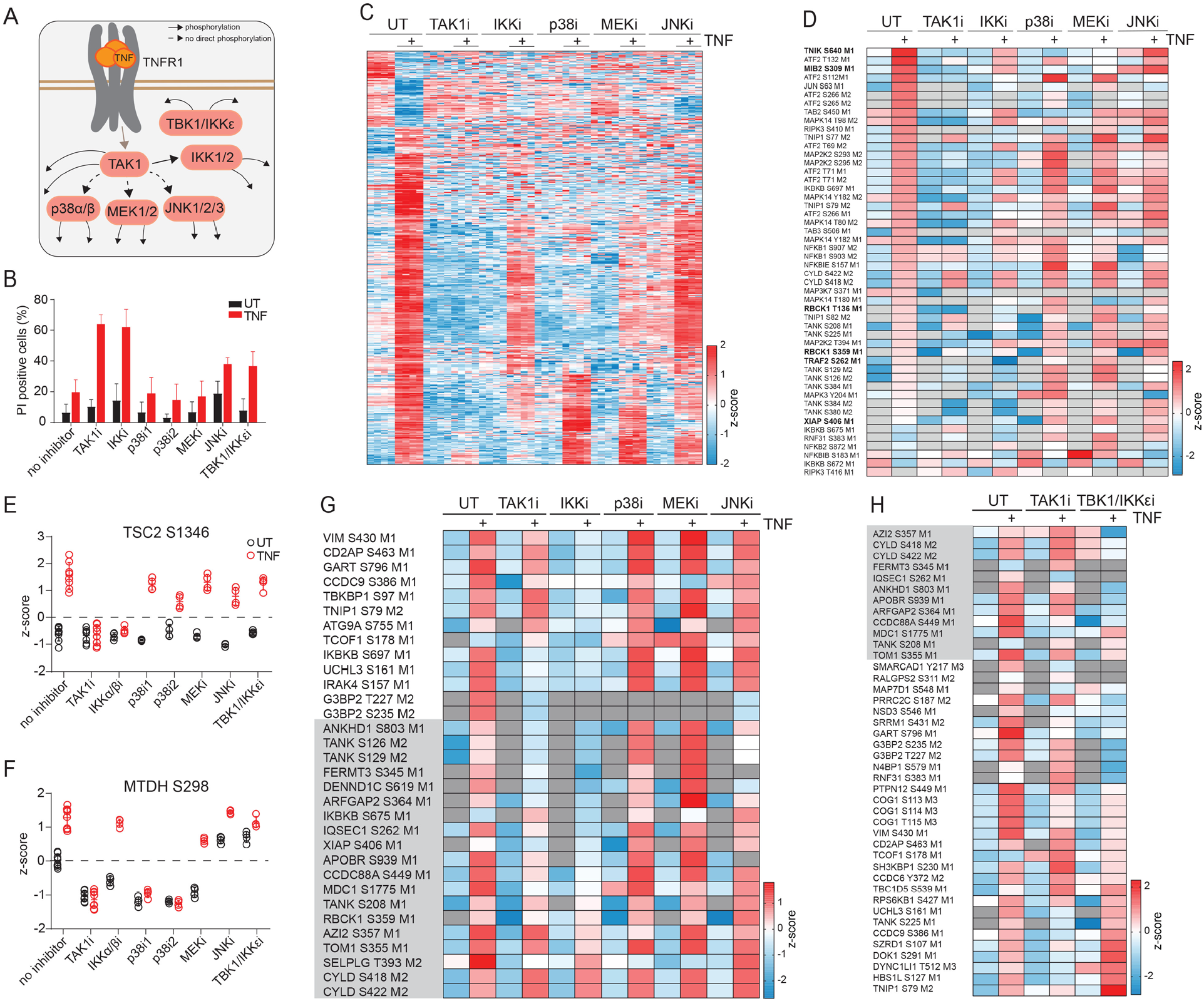
Kinase inhibition unravels kinase-substrate relations upon TNF stimulation. (**A**) TNF signaling scheme highlighting kinase hubs that we inhibited in this study. (**B**) Cell death analysis by flow cytometry of propidium iodide-stained U937 cells either stimulated with the indicated kinase inhibitors alone (TAK1 inhibitor (7-Oxozeaenol, 1 μM), IKK inhibitor (TCPA-1, 5 μM), p38 inhibitor 1 and 2 (LY228820, 2 μM; SB203580, 10 μM), MEK inhibitor (PD025901, 10 μM), JNK inhibitor (SP600125, 20 μM)) or pretreated with kinase inhibitors 1 h before TNF treatment for 24 h (± SD, n ≥ 2). (**C**) Heatmap of z-scored phosphosite intensities significantly changing in U937 cells treated for 15 min with TNF compared to untreated cells (Student’s t-test FDR < 0.05). Cells were additionally pretreated for 1 h with kinase inhibitors as indicated in (**B**). Z-scores of the four replicates are shown. (**D**) Heatmap of means of z-scored phosphosite intensities on known classical members of the TNF signaling pathway. U937 cells were treated as in (**C**). (**E, F**) Z-scored phosphorylation levels of S1346 on TSC2 (**E**) and S298 on MTDH (**F**) in untreated (open black circles) and TNF-treated cells (red open circles) in the presence of the indicated kinase inhibitors (± SD). (**G**) Heatmap of means of z-scored phosphosite intensities that are induced upon TNF stimulation despite TAK1 inhibition (Student’s t-test of cells treated with TAK1 inhibitor alone compared to TAK1 inhibitor with TNF, -Log10 p >1.3, Log2 fold change > 1.3 using two different imputations, **Methods**). Phosphosites with grey background are not ablated by any inhibitor used. (**H**) Heatmap of means of z-scored phosphosite intensities that are induced upon TNF stimulation despite TAK1 inhibition (Student’s t-test of cells treated with TAK1 inhibitor alone compared to TAK1 inhibitor with TNF, -Log10 p >1.3, Log2 fold change > 1.3 using two different imputations). Cells were treated with: TAK1 inhibitor (7-Oxozeaenol, 1 μM) and TBK1/IKKε inhibitor (MRT67307, 2 μM). Phosphosites with grey background were not ablated by any inhibitor used in the experiment of (**G**).

These experiments also identified upstream kinases responsible for phosphorylation events on classical TNF signaling complex members (**Figure 2D**). Phosphorylation of the TRAF2 and NCK-interacting protein kinase (TNIK) (S640) and of Mind bomb-2 (MIB2) (S309), a newly discovered E3 ligase in TNF signaling (Feltham et al., 2018), was inhibited by TAK1, p38, and MEK inhibition. XIAP and TRAF2 phosphorylation was strongly reduced by TAK1 and IKK2 inhibition. To illustrate the richness of our data, we note that our data confirm a previous report that showed that IKK2 phosphorylates Tuberous Sclerosis Complex 2 (TSC2) at position S1346, which triggers mTOR signaling downstream of the TNF receptor (Lee et al., 2007) (**Figure 2E**). A recent phosphoproteomics study reported that the phosphorylation of Metadherin (MTDH) at position S298 is important for NF-κB signaling and is also mediated by IKK2 (Krishnan et al., 2015). However, we failed to detect downregulation of this site due to the IKK1/2 inhibitor, but instead measured inhibition with two different p38 inhibitors in two independent experiments (**Figure 2F**). Surprisingly, phosphorylation sites on TNFAIP3 interacting protein 1 (TNIP1), TANK-binding kinase 1-binding protein 1 (TBKBP1), Autophagy-related protein 9A (ATG9A), and other proteins were not abrogated by TAK1 inhibition but were strongly affected by IKK1/2 inhibition (**Figure 2G**). This demonstrates that IKK1/2 must retain some function independent of TAK1 activity.

Furthermore, we detected two novel phosphorylation sites (T136 and S359) on HOIL1, also called RBCK1 for RanBP-type and C3HC4-type zinc finger-containing protein 1, an essential member of the Linear ubiquitin assembly complex (LUBAC) (**Figure 2D**). The phosphorylation at T136 was inhibited by blocking TAK1 and IKK2 activity, while phosphorylation at S359 did not change significantly. Phosphorylation sites on other proteins associated with LUBAC, including the Ubiquitin carboxyl-terminal hydrolase (CYLD) and the 5-azacytidine-induced protein 2 (AZI2) also occurred independently of the kinase inhibitors used above (**Figure 2G**). LUBAC is required for the recruitment and activation of TANK-binding kinase 1 (TBK1)/ Inhibitor of nuclear factor kappa-B kinase subunit epsilon (IKKε), and IKKε phosphorylates and activates the deubiquitinase CYLD (Hutti et al., 2009, Lafont et al., 2018). Therefore, we tested whether TBK1 and IKKε are responsible for phosphorylating other LUBAC members and associated proteins. Indeed, inhibiting TBK1/IKKε affected not only TNF-induced phosphorylation of CYLD but most of the other proteins whereas TAK1 inhibition failed to block these phosphorylation events, including on ADP-ribosylation factor GTPase activating protein 2 (ARFGAP2), IQ motif and SEC7 domain-containing protein 1 (IQSEC1), Coiled-coil domain containing 88b (CCDC88A), and Fermitin family homolog 3 (FERMT3) (**Figure 2H**). Our data thus suggests a potential association of these proteins with the LUBAC complex upon TNF signaling.

### TNF-mediated phosphorylation induces wide-spread protein re-localization

TNF-dependent phosphorylation relies on the correct localization of kinases and their substrates and requires subcellular protein translocation (Micheau and Tschopp, 2003). TNF binding to its receptor recruits a range of complex members to the plasma membrane and binding of TRAF2, and requirement of TRADD, RIPK1, IAPs, and the LUBAC complex for proper activation of TAK1 is well documented (Brenner and Shenderova, 2015). However, the impact of phosphorylation on protein localization is less explored. We therefore set out to obtain a systems-based view on protein translocations and the role of phosphorylation on these translocations upon TNF stimulation. We treated cells with TNF for 15 min with and without the TAK1 inhibitor and subsequently separated membrane, nucleus, and cytosol (**Figure 3A**). Peptides were either phospho-enriched or directly measured for full proteome analysis. Compartment markers were strongly enriched in their respective fraction, indicating successful fractionation (**Supplementary Figure S3A**). Most measured proteins and phosphopeptides were significantly enriched in one of the indicated fractions (**Figure 3B, C**).

**Figure 3:**
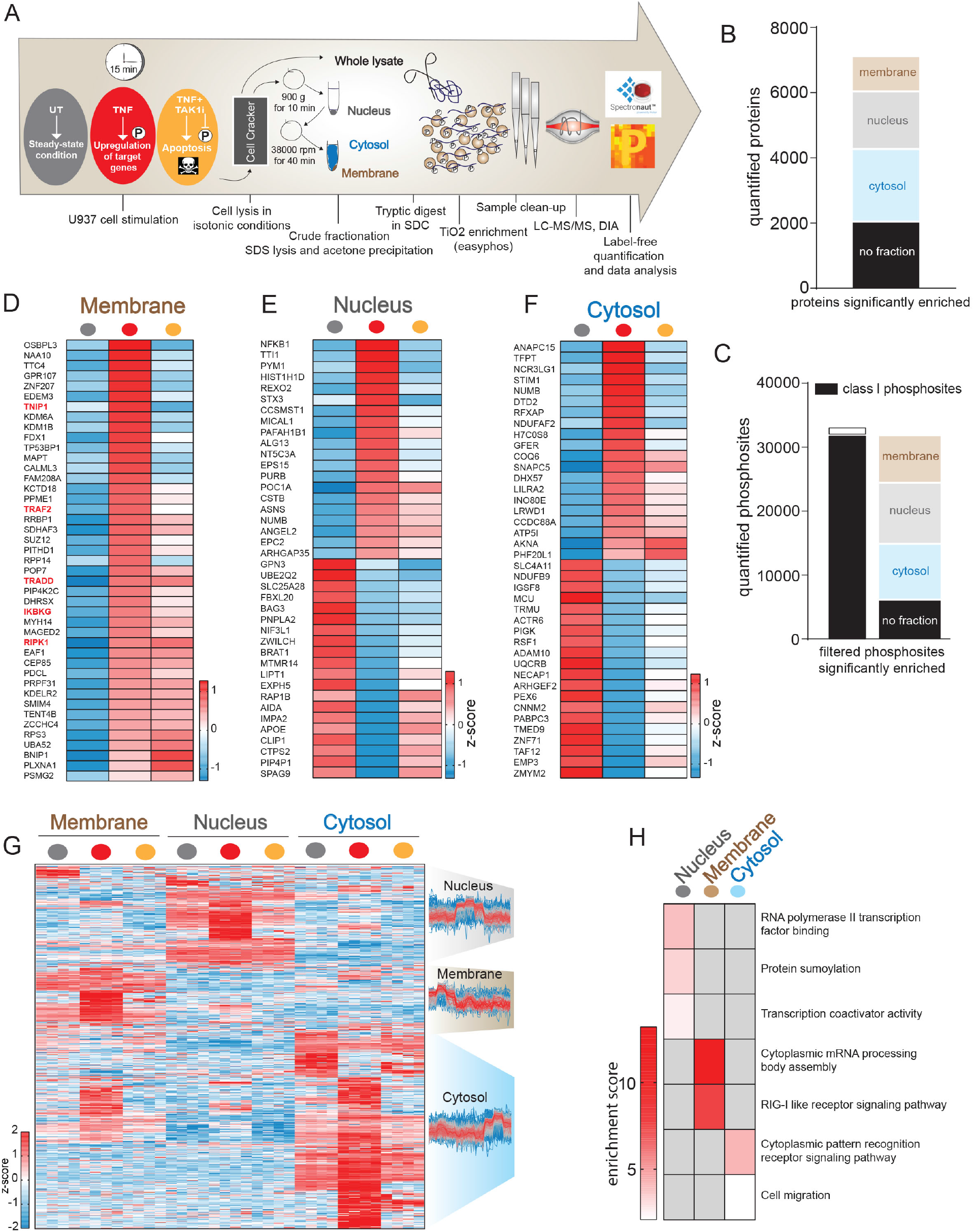
TNF-induced phosphorylation strongly affects translocation of signaling members downstream of TAK1. (**A**) Workflow of crude fractionation with subsequent proteome and phosphoproteome analysis of U937 cells. Grey = untreated, red = TNF, orange = TNF+TAK1 inhibitor (**B**) Most quantified proteins are significantly enriched in one of membrane, nucleus, and cytosol fractions (Student’s t-test FDR < 0.05). (**C**) Total number of quantified phosphosites and the numbers of phosphosites significantly enriched in the three fractions (Student’s t-test FDR < 0.05). Class I sites (in black) are phosphosites with a localization probability of > 75%. (**D**) Heatmap of means of z-scored protein intensities enriched in the membrane upon TNF treatment (Student’s t-test, -Log10 p > 1.29, Log2 fold change > 0.5). (**E**) Heatmap of means of the 20 most strongly up- and downregulated z-scored protein intensities in the nucleus upon TNF treatment (Student’s t-test -Log10 p > 1.5, Log2 fold change > 0.5 or < -0.5). (**F**) Heatmap of means of the 20 most strongly up- and downregulated z-scored protein intensities in the cytosol upon TNF treatment (Student’s t-test -Log10 p > 1.5, Log2 fold change > 3 or < -3). (**G**) Heatmap of z-scored phosphosite intensities that are significantly regulated upon TNF treatment in the membrane, nucleus, and cytosol (Student’s t-test FDR < 0.05). The profiles are color coded according to their distance from the respective cluster center (red is close to center, blue is further away from center). (**H**) Fisher’s exact test of phosphosites significantly enriched in nucleus, membrane (p < 0.001) and cytosol (p < 0.01). The red scale represents enrichment, while grey represents no enrichment.

We detected TRADD, RIPK1, and TRAF2 at the plasma membrane after stimulation, providing a positive control of our spatial proteomics experiment (**Figure 3D**). Their recruitment is TAK1 independent, confirming their upstream role in TNF signaling. NF-κB essential modulator (NEMO or IKBKG) and TNIP1 levels also increased in the membrane fraction upon TNF stimulation (**Figure 3D**). While S77, S82 and S79 on TNIP1 were independent of TAK1 activity (**Figure 2D, G**), its recruitment to the membrane was TAK1 dependent (**Figure 3D**). Apart from known TNF-complex members, several other proteins also translocated to the membrane fraction. Although this does not prove their direct association with the TNF receptor complex, it demonstrates their translocation during TNF signaling. Some proteins dissociated from the membrane compartment upon TNF treatment (**Supplementary Figure 3B**). The most regulated proteins at the membrane were enriched for proteins involved in the death domain-mediated complex assembly and the regulation of necrotic cell death (p < 0.002; Fisher’s exact test; **Supplementary Figure 3C**). Surprisingly we detected increased levels of TAK1 (MAP3K7), IKK1 (CHUK), and TAB1/2 in the membrane upon TAK1 inhibition (**Supplementary Figure 3D)**. This may be due to a stabilizing effect of TAK1 inhibitor on TAK1 at the membrane.

Within 15 min of stimulation, TNF triggered nuclear translocation of many proteins involved in transcription, indicating a transcriptional response (**Figure 3E**). The transcription factor NFΚB1 was the most strongly enriched protein in the nucleus and this was entirely prevented by TAK1 inhibition (**Figure 3E**). TAK1 inhibition also increased RIPK1 levels in the nucleus, suggesting that phosphorylation of RIPK1 prevents its nuclear translocation (**Supplementary Figure 3E**). Protein translocation to and from the cytosol is also heavily phosphorylation dependent (**Figure 3F**).

Most kinases, including TAK1 (MAP3K7), p38 (MAPK14), IKK2 (IKBKB), MEK1/2 (MAP2K1/2), are primarily enriched in the cytosol, independent of TNF stimulation (**Supplementary Figure 3F**). TNF-regulated phosphosites on proteins downstream of these kinases are, however, enriched in all cellular compartments (**Figure 3G**). For example, in the nucleus we detected TNF-regulated phosphosites on proteins involved in RNA polymerase II transcription (**Figure 3H**). Proteins of the RIG-I pathway and cytoplasmic RNA processing body assembly are differentially phosphorylated in the membrane, and phosphosites within pattern recognition receptor signaling and cell migration pathways are regulated in the cytosol (**Figure 3H**). Most of these are dependent on the activity of TAK1, which is localized in the cytosol, implying that TNF triggers protein translocation of kinases and their substrates (**Figure 3G**). We even measured TNF-regulated phosphorylation events on peptides within fractions where we failed to detect the total protein. This indicates that phosphorylations of these substrates trigger translocation, or that upon translocation, the proteins are phosphorylated immediately (**Supplementary Figure 3G-J**).

### TNF-induced cell death triggers a strong RNA-processing response and CDK activation

Inhibition of the NF-κB signaling pathway by targeting TAK1 and IKK2 induces cell death downstream of TNF (**Figure 2B**) (Mihaly et al., 2014). This cell death can either be apoptotic (caspases active) or necroptotic (caspases inactive). To dissect the role of phosphorylation events regulated upon cell death, we treated U937 cells for three hours with TNF alone, or in combination with the cIAP inhibitor Smac-mimetic (Birinapant) to induce apoptosis, or with Smac-mimetic and the caspase inhibitor Idun (IDN-6556) to induce necroptosis (**Figure 4A**). We also performed experiments in BMDM cells that were pretreated with TAK1 and caspase inhibitors prior to TNF stimulation **(Supplementary Figure 4A)**. This identified several phosphorylations on members of the TNF signaling pathway (**Figure 4A, B**). The classical activating phosphorylations S418/S422 on CYLD were reduced, more likely due to CYLD cleavage by active caspases than active dephosphorylation (O’Donnell et al., 2011) (**Figure 4B**). S5, S406 and S430on XIAP, an E3 ligase known to inhibit TNF-induced cell death, were also strongly regulated upon caspase activation (**Figure 4B**). S430 induces XIAP auto-ubiquitination, resulting in its degradation, leading to increased cell death upon viral infection (Nakhaei et al., 2012). However, when we tested the impact of the phosphomimetic and -ablating XIAP mutants on TNF-induced cell death, we failed to detect differences compared to cells expressing XIAP wildtype (**Supplementary Figure 4B, C**). The impact of these phosphorylations might depend on the cell system used. In BMDMs, we detected RIPK3 phosphosites that were upregulated during necroptosis (**Supplementary Figure 4D, E**). S14 and S25 of RIPK1 similarly increased during cell death, whereas S15, S313, S321, and S415 did not. With respect to major cellular processes regulated during apoptosis, we found phosphosites on proteins involved in RNA splicing and mRNA processing significantly upregulated in both cell types (344 phosphosites in U937 cells and 123 phosphosites in BMDMs, FDR < 0.05 Student’s t test) (**Figure 4C; Supplementary Figure 4F, G, H**). These processes were not only induced upon caspase activation but to a smaller degree also during necroptosis (**Figure 4C; Supplementary Figure 4G)**. In contrast, the activation of the DNA damage sensor kinase Ataxia telangiectasia mutated (ATM) upon TNF-induced cell death was strictly dependent on caspase activation. Indeed, DNA fragmentation via the caspase-activated DNase triggers this DNA damage response upon TNF-induced apoptosis (McIlroy et al., 1999) (**Figure 4C; Supplementary Figure 4F, H**). Necroptosis induced the phosphorylation of membrane proteins, most likely as a consequence of plasma membrane permeabilization in U937 cells, and it caused a strong GTPase response in BMDMs (**Figure 4C; Supplementary Figure 4F**). Proteome analyses in both cell lines revealed that stimulation-induced phosphoproteome changes did not correlate with proteome changes (**Supplementary Figure 4I-L**).

**Figure 4:**
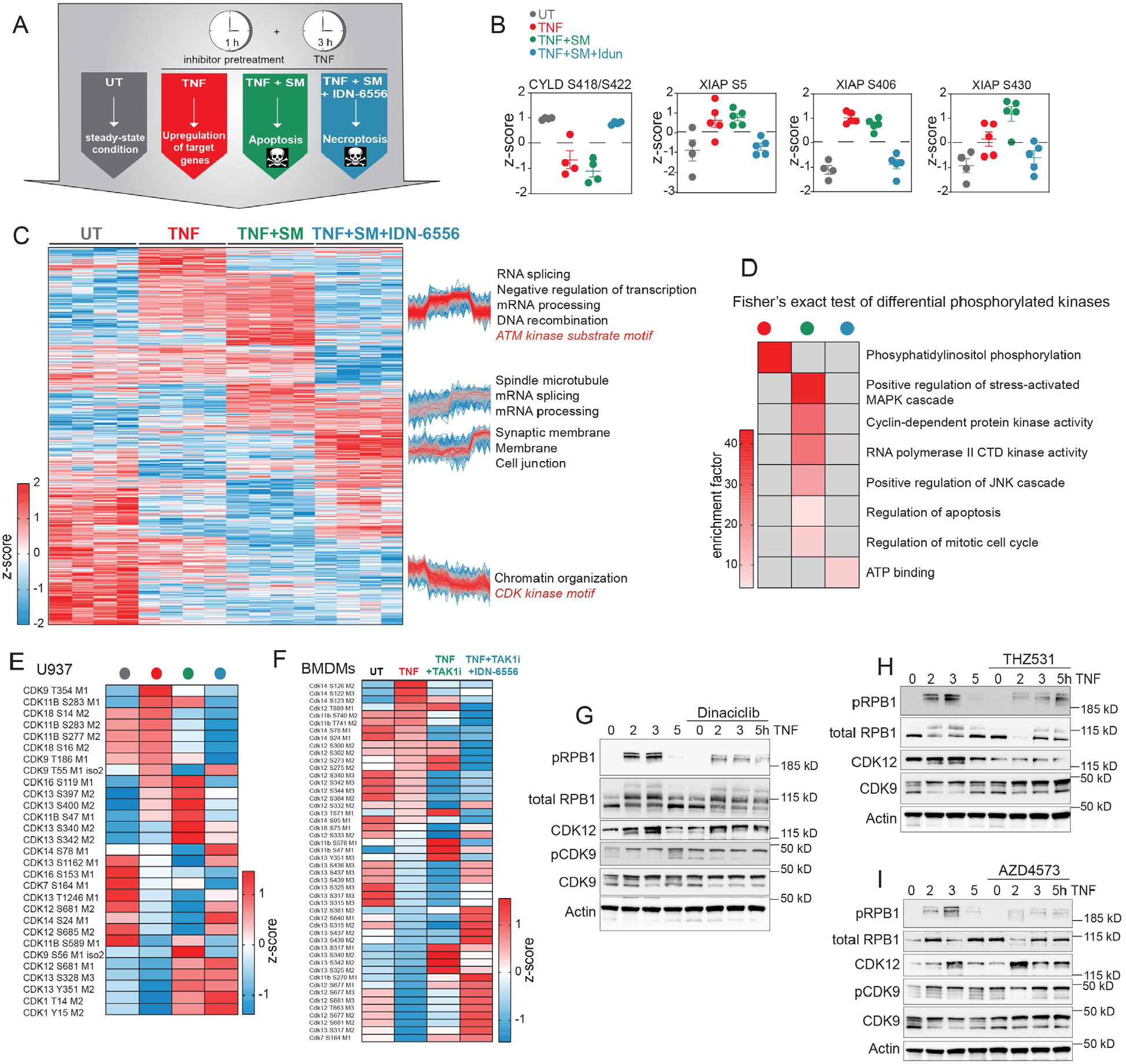
TNF-induced cell death leads to a strong RNA processing response and regulation of the phosphorylation status of CDKs. (**A**) Experimental scheme of U937 cells that were untreated, treated with TNF (100 ng/ml, red) alone, treated with TNF and Smac-mimetic (SM/Birinapant, 1.25 μM, green) to induce apoptosis or with TNF, SM and IDN-6556 (Emricasan, 10 μM, blue) to induce necroptosis. (**B**) Z-scored phosphosite intensities of CYLD and XIAP that are regulated upon TNF treatment, TNF-induced apoptosis, and TNF-induced necroptosis (± SD). XIAP phosphosites were retrieved from a DDA dataset (Methods). (**C**) Heatmap of z-scored phosphosite intensities that are significantly ANOVA regulated upon TNF treatment or TNF-induced cell death (FDR < 0.05). The profiles are color coded according to their distance from the respective cluster center (red is close to center, blue is further away from center). (**D**) Fisher’s exact test on kinases with significantly regulated phosphosites upon different treatments (FDR < 0.02). (**E**) Heatmap of means of z-scored phosphosite intensities of CDKs that significantly changed (ANOVA) upon treatment of U937 cells (FDR < 0.05). (**F**) Heatmap of means of z-scored phosphosite intensities of CDKs that changed significantly (ANOVA) upon treatment of BMDMs with TNF (red) alone, TNF and TAK1 inhibitor (TAKi) to induce apoptosis (green) or TNF, TAK1i and the caspase inhibitor IDN-6556 to inducenecroptosis (blue). Untreated cells (UT) served as controls (FDR < 0.05).(**G-I**) Immunoblots of U937 cells stimulatedwith TNF as indicated or in combinationwith CDK inhibitors Dinaciclib (6 nM)(**G**), THZ531 (200 nM)(**H**), AZD4573 (6 nM)(**I**) and probed with antibodies against phosphorylated (S2) and total (N-terminal) RPB1,phosphorylated (T186) and total CDK9, CDK12 and β-Actin (n = 2).

Phosphorylation of kinase motifs in CDK1, 4, 5, and 6 in apoptotic and necroptotic cells were downregulated, suggesting inhibition of the cell cycle during cell death (**Figure 4C; Supplementary Figure 4M**). Enrichment analysis on kinases whose phosphorylation was regulated upon stimulation revealed regulation of cyclin-dependent protein kinases during apoptosis (**Figure 4D**). Filtering for ANOVA significantly changing phosphosites on CDKs upon stimulation, showed that transcriptional CDKs (CDK7, 9, 12, 13, and 14) were regulated rather than CDKs involved in the cell cycle regulation (Methods, **Figure 4E, F**). Transcriptional CDKs modulate transcription primarily by regulating RNA polymerase II activity (Malumbres, 2014) (**Figure 4D**). This occurs by phosphorylation of the carboxy-terminal domain (CTD) of its largest subunit RPB1, which consists of 52 heptad sequences (**Supplementary Figure 4N**). CDK9 is part of the p-TEFb complex, which induces RPB1 mediated transcriptional elongation by phosphorylating its CTD repeats (Price, 2000). Following three hours of TNF treatment phosphorylation on T354 of CDK9 increased 9-fold (**Figure 4E**). This C-terminal site is autophosphorylated by active CDK9 (Garber et al., 2000). To demonstrate CDK9 activation upon TNF stimulation, we blotted for the CDK9 activating phosphorylation at position T186 and observed a moderate upregulation (**Figure 4G, I**). Many phosphorylation sites on both CDK12 and 13 were differentially regulated upon TNF treatment and TNF-induced cell death (**Figure 4E, F**). The role of CDK12 and CDK13 in RPB1-mediated transcription was discovered a decade ago, and more recently, CDK12 was shown to be important for intact expression of long genes and reported as a potential anti-tumor target (Greenleaf, 2019, Krajewska et al., 2019). However, little is known about their regulation via phosphorylation. To test whether these transcriptional CDKs actually phosphorylate RNA polymerase II upon TNF treatment, we blotted for phosphorylation of RPB1 at the CTD heptad sequence position S2. We detected increased phosphorylation of RPB1 upon TNF stimulation, which was reduced by the pan-CDK inhibitor Dinaciclib (**Figure 4G-I**). Importantly, THZ531 and AZD4573, specific inhibitors of CDK12/13 and CDK9, respectively (Zhang et al., 2016, Cidado et al., 2020, Krajewska et al., 2019), also inhibited phosphorylation of S2 of RPB1, with AZD4573 exhibiting a more potent effect (**Figure 4H, I**). This indicates that TNF signaling induces phosphorylation and thereby CDK9 and CDK12/13-mediated activation of RPB1.

### CDK kinase activity is required for the transcription of TNF target genes

TNF triggers a potent transcriptional response by inducing a wide range of target genes. Its proinflammatory properties are primarily mediated through the upregulation of cytokines, which are required to fight infections, while the upregulation of pro-survival genes prevents excessive cell death and inflammation. As our data demonstrated regulated RPB1 phosphorylation and as this protein is required for TNF-mediated transcription, we wondered if inhibition of transcriptional CDKs would impact TNF-induced target gene expression. Global phosphoproteomics of CDK inhibitor-treated cells inhibited TNF-induced phosphorylation of CDK12/13 as well as RNA processing at early and late time points (**Supplementary Figure 5A, B, C**). Interestingly, proteome analysis of TNF-treated U937 cells revealed that Dinaciclib or the two CDK12 inhibitors CDK12-IN3 (Johannes et al., 2018) and THZ531 almost completely ablated protein regulation (**Figure 5A**). These CDK inhibitors also decreased upregulation of the classical target genes ICAM1 and SOD2, albeit not to the same extent (**Figure 5B**). Likewise, there was less upregulation of OTULIN, a DUB that deubiquitylates LUBAC and thereby prevents cell death (Heger et al., 2018). The slight downregulation of XIAP was abrogated by CDK inhibition, which indicates that this regulation is dependent on transcription (**Figure 5B**). NF-κB signaling itself – the main pathway driving TNF-induced transcription - was unaffected **(Figure 5C, Supplementary Figure 5D)**. To verify that CDK inhibitors indeed inhibit the expression of target genes at the transcriptional level, we examined their impact on TNF-mediated upregulation of MCP-1 mRNA levels, which were indeed reduced (**Figure 5D**). Similarly, less of the cytokine IP10 was released from BMDMs upon addition of CDK inhibitors (**Supplementary Figure 5E**). Consistent with previous studies that investigated the role of CDK9, this indicates that inhibiting transcriptional CDKs 9, 12, and 13 may have an anti-inflammatory effect in disease (Krystof et al., 2012, Henry et al., 2018). The induction of the pro-survival protein FLIP was also strongly inhibited by CDK inhibitors (**Figure 5E-H**). Pan-CDK and CDK9 inhibition strongly reduced FLIP_L_ and FLIP_S_ expression upon TNF stimulation (Figure 5E, F), while CDK12/13 inhibitors reduced FLIP_L_ but increased FLIP_S_ levels compared to TNF stimulation alone (**Figure 5G, H; Supplementary Figure 5F**).

**Figure 5:**
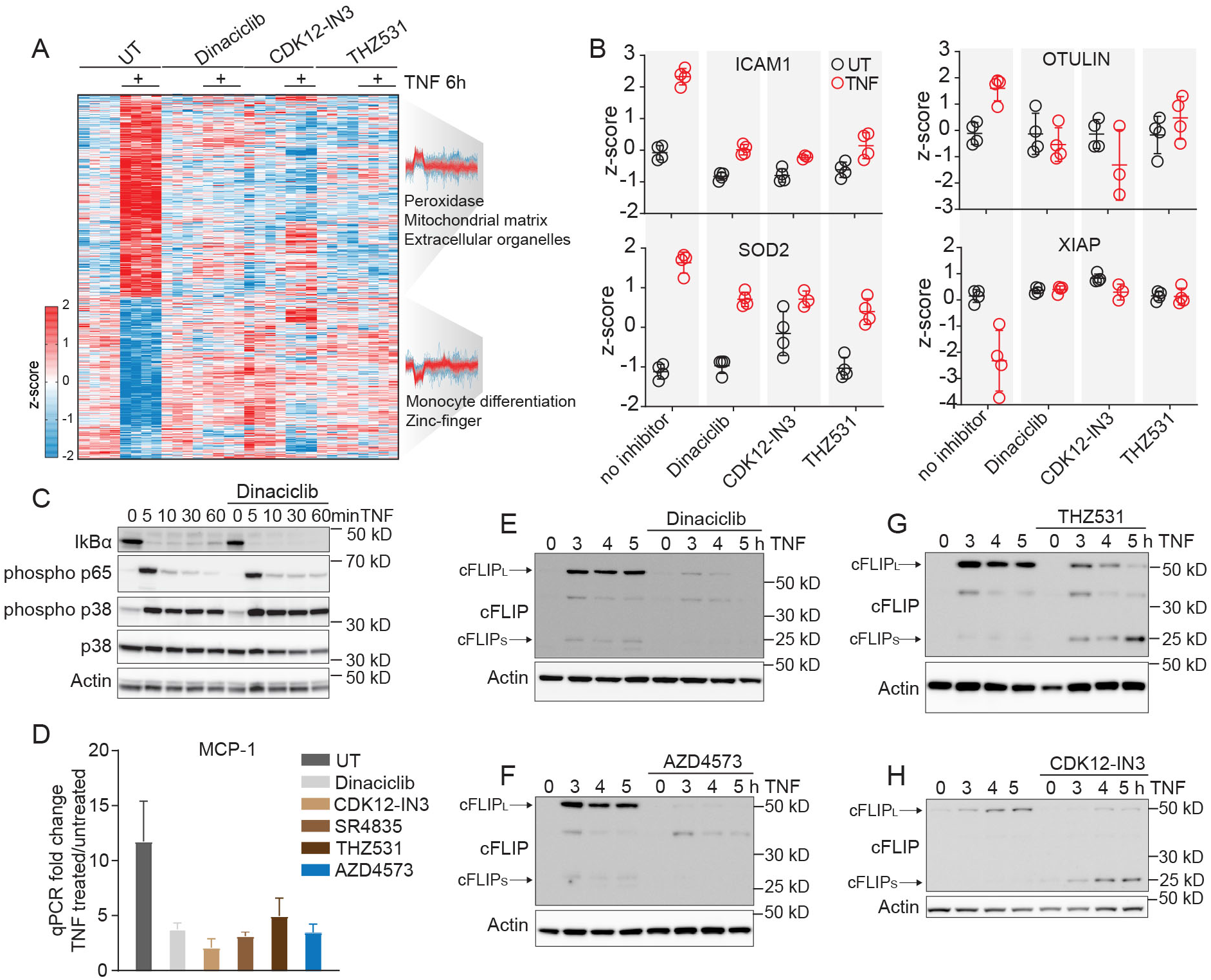
CDK9 and CDK12/13 inhibitors potently reduce transcription of TNF-target genes. (**A**) Heatmap of z-scored protein intensities significantly changed upon TNF treatment for 6 h in U937 cells (FDR < 0.05). Cells were additionally treated with Dinaciclib (6 nM), CDK12-IN3 (60 nM), and THZ531 (400 nM). Fisher’s exact test on up- and downregulated protein clusters (Student’s t-test, FDR < 0.001). The profiles are color coded according to their distance from the respective cluster center (red is close to center, blue is further away from center). (**B**) Z-scored protein levels of selected TNF target genes and members of the TNF signaling pathway. (**C**) Immunoblot of U937 cells treated with TNF alone and in combination with the pan-CDK inhibitor Dinaciclib. Proteins were blotted for IκBα, phosphorylated p65, phosphorylated and, total p38 and β-Actin (loading control). (**D**) qPCR of MCP1 of U937 cells treated for 4 h with TNF alone or in combination with CDK inhibitors (n = 3). (**E-H**) U937 cells treated with TNF alone and in combination with CDK inhibitors were blotted for cFLIP and β-Actin (n = 2).

### Transcriptional CDKs inhibit TNF-induced cell death

FLIP inhibits apoptosis through binding to caspase-8, thereby preventing its activation (Wilson et al., 2009). FLIP/caspase-8 heterodimers also potently inhibit necroptosis (Oberst et al., 2011). A previous study presented CDK9 as a potential target for tumor therapy in combination with TRAIL treatment (Lemke et al., 2014). Therefore, we wondered whether inhibition of TNF-mediated FLIP upregulation by transcriptional CDK inhibitors would enhance TNF-induced cell death. Treating cells with the pan-CDK inhibitor dinaciclib, the CDK9 inhibitors Flavopiridol and AZD4573, the CDK12 inhibitor CDK12-IN3 or the CDK12/13 inhibitors THZ531 and SR4835 all triggered synergistic cell death in combination with TNF, TNF and Smac-mimetic (SM) or SM and IDN-6556 (**Figure 6A-D; Supplementary Figure 6A, B**). Inhibitors targeting CDKs that are not involved in transcription failed to induce synergistic cell death, while CDK12 knockdown moderately increased TNF-dependent cell death (**Supplementary Figure 6A, C, D**). These experiments show that transcriptional, but not non-transcriptional CDKs exacerbate TNF induced cell death.

**Figure 6:**
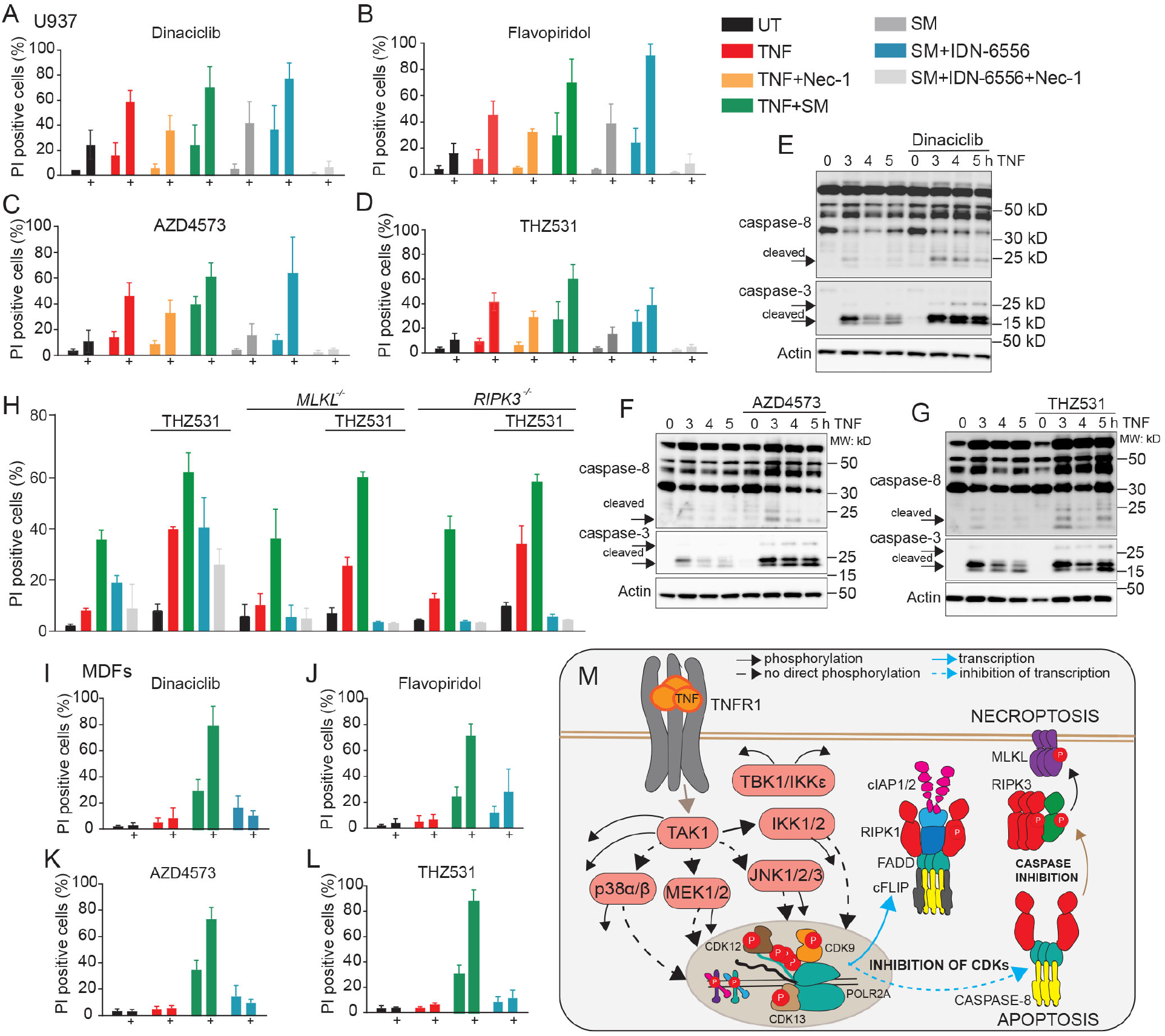
CDK9 and CDK12/13 inhibit TNF-induced cell death. (**A-D**) Cell death analysis by flow cytometry of propidium iodide-stained U937 cells. Cells were pretreated for one h with CDK inhibitors Dinaciclib (6 nM) (**A**), Flavopiridol (60 nM) (**B**), AZD4573 (6 nM), (**C**) THZ531 (200 nM) (**D**) or left untreated (black) before treatment with TNF (red), TNF and Necrostatin-1 (50 μM, yellow), TNF and SM (Birinapant, 1.25 μM, green), SM alone (grey), SM and IDN-6556 (10 μM, blue) or SM, IDN6556 and Necrostatin-1 (light grey) for 24 h (± SD, n ≥ 3). (**E-G**) Immunoblots of U937 cells stimulated with TNF or TNF in combination with CDK inhibitors stained for caspase-3 and 8 and β-Actin (n = 2). (**H**) Cell death analysis by flow cytometry of propidium iodide-stained wt and MLKL and RIPK3 deficient U937 cells stimulated as described for 24 h (± SD, n ≥ 3). (**I-L**) Flow cytometry analysis of propidium iodide stained wildtype MDFs treated as indicated (SM, Compound A (2 μM); pan-CDK inhibitor, Dinaciclib (12 nM); CDK9 inhibitors, Flavopiridol (60 nM), AZD4573 (6 nM); CDK12/13 inhibitors, THZ531 (800 nM)) (± SD, n ≥ 3). (**M**) TNF signaling scheme highlighting the role of the kinases and specifically CDKs in cell death as inferred from our data.

CDK9 and CDK12/13 inhibition, in combination with TNF stimulation, enhanced apoptosis, as evidenced by increased caspase-8 and caspase-3 cleavage (**Figure 6E-G, Supplementary Figure 6E**). This only depended slightly on RIPK1 kinase activity (**Figure 6A-D; Supplementary Figure 6B**). Inhibition of cIAP1/2 by SM alone - like CDK inhibition - had a minor impact on cell death, which was increased by their combination, and especially by the addition of TNF (**Figure 6A-D; Supplementary Figure 6B**).

To test if CDK inhibitors also enhance TNF-dependent necroptosis, we treated U937 cells with a combination of SM and the caspase inhibitor IDN-6556 (Brumatti et al., 2016). This showed that CDK inhibition also enhanced necroptosis in wildtype U937 cells, whereas cells lacking the necroptotic mediators MLKL and RIPK3 failed to die (**Figure 6A-D, H; Supplementary Figure 6C, F, G**). This synergistic cell death is not restricted to U937 cells but also present in mouse dermal fibroblasts (MDFs) and murine BMDMs (**Figure 6I-L; Supplementary Figure 6H, I**). Unlike pan-CDK inhibitors, CDK12 specific inhibitors failed to induce synergistic cell death in BMDMs, most likely due to sustained levels of FLIP_L_ and the upregulation of FLIP_S_ (**Supplementary Figure 6I, J**). TNF alone was not sufficient to induce synergistic cell death in these cells, suggesting that CDK inhibition can only potentiate but not trigger cell death (**Figure 6I-L; Supplementary Figure 6 H, I**). We conclude that TNF-mediated regulation of transcriptional CDKs results in phosphorylation and regulation of their substrates including the CTD of RPB1. Because this largest RNA Polymerase II subunit is necessary for transcription of pro-survival proteins like FLIP, its phosphorylation is in turn necessary to prevent apoptosis and necroptosis (**Figure 6M**).

## Discussion

TNF primarily exerts its functions through the transcriptional regulation of multiple target genes, which is strongly dependent on the integration of a number of different signaling events. Furthermore, exact timing of signaling events is essential for appropriate signaling. Here, we aimed to elucidate the kinetics of TNF-induced phosphorylation events on a global scale. Our in-depth, quantitative phosphoproteome analysis revealed that early and late TNF signaling is characterized by distinct phosphorylation events, which broadly represented upstream and downstream signaling events.

Importantly, TNF stimulation never occurs in isolation and upon infections TLRs and inflammasomes are activated, leading to the production of a range of cytokines. Indeed, our data showed that TNF also induced a robust and dynamic regulation of phosphorylation on proteins involved in other immune signaling pathways. We observed TNF-induced phosphorylation events of the inflammasome components like NLRC4, adaptor proteins like MAVS, kinases like IRAK4, and phosphatases like INPP5D and speculate that they could act to prime and interlink immune signaling pathways to either enhance or dampen signaling strength. Our data suggest that this cross-talk is most likely mediated through the activation of mutual kinases, congruent with the fact that TAK1, IKKs, p38, MEK1,2, and JNK, for example, are activated by various immune ligands. Inhibition of the TAK1 showed that the vast majority, but not all TNF-regulated phosphorylation events are dependent on this master kinase. The few TAK1-independent phosphosites were instead regulated by IKK2, TBK1, or IKKε. It has recently been shown that TBK1 and IKKε are recruited to the TNF signaling complex (Lafont et al., 2018) where IKKε phosphorylates CYLD. Here we have identified several other likely substrates of TBK1 and IKKε, including members of the LUBAC and speculate that they localize to LUBAC’s vicinity.

To identify kinases apart from IKKs and MAPKs that act downstream of TAK1 and contribute to TNF-mediated cytokine production and cell death, we analyzed the activation of kinases upon TNF-induced apoptosis and necroptosis. This uncovered an extensive phospho-regulation of transcriptional CDKs, including CDK9, 12, and 13. Unlike the inhibition of TAK1 or IKK, CDK inhibition alone was not sufficient to trigger cell death upon TNF treatment. CDK9 and CDK12/13 inhibition could have a weaker impact on TNF-induced transcription compared to TAK1 and IKK inhibition or additional functions of these latter kinases on the TNF signaling complex may directly contribute to cell death induction independent of transcription.

CDK9 and CDK12/13 inhibitors are currently tested for their efficiency to kill cancer cells (Galbraith et al., 2019). In particular, CDK12, which also plays a role in RNA processing/splicing and upregulation of DNA repair genes (Dubbury et al., 2018), has recently been reported to frequently be mutated in cancer, making it a promising cancer therapy target (Quereda et al., 2019). Here we show that their activity strongly modulates the upregulation of most TNF-induced genes, including the caspase-8 inhibitor protein cFLIP. Reduction in CDK activity or levels induced synergistic cell death with TNF in transformed cells. Tumor therapy might profit from our finding that not only CDK9 (Lemke et al., 2014) but also CDK12/13 inhibitor-mediated killing of cancer cells can be enhanced by cell death-inducing cytokines and ligands. Furthermore, CDK inhibition could also be useful in anti-viral therapy. Viruses such as HIV exploit CDKs for viral transcription (Salerno et al., 2007). Therefore, targeting CDKs may both inhibit the transcription of viral genes and also enhance cell death of infected cells.

Cytokine expression induced by TNF was also heavily affected by CDK9 and CDK12/13 inhibitors. Previous studies already proposed targeting CDK9 as an anti-inflammatory treatment (Schmerwitz et al., 2011). Here we demonstrated that CDK12/13 inhibition also potently impacts cytokine upregulation by TNF. Although transcriptional CDK inhibitors reduced the expression of proinflammatory cytokines, they increased TNF-induced cell death substantially, which in turn exacerbates inflammation. This might explain increased levels of IL-6 in the pan-CDK inhibitor Flavopiridol-treated patients suffering from proinflammatory syndrome (Messmann et al., 2003). In contrast to the CDK9 and pan-CDK inhibitors, CDK12 inhibition upregulated FLIPs levels. Our observation that CDK12 inhibition failed to induce synergistic cell death in BMDMs suggests this protein as a potentially more suitable target for anti-inflammatory therapy.

In conclusion, our in-depth phosphoproteome analysis of TNF-stimulated myeloid cells elucidated the kinetics, kinase-substrate relations, and localizations of thousands of phosphorylation event. In a cell death setting, we detected strong regulation of transcriptional CDK phosphorylation, which we subsequently identified to be necessary for intact RPB1 phosphorylation and transcription of TNF-target genes. We propose that depending on the context, inhibitors against different transcriptional CDKs could be used as anti-inflammatory agents or tumor cell killers. Our work provides a valuable resource of TNF-regulated phosphorylation events to the community. As importantly, it discovers an aspect of TNF signaling that is essential for intact transcription of TNF-target genes, thereby contributing to the decision between cell death and survival, auto-inflammation, or an efficient response to infections.

## Supporting information

TNF 15 min BMDMs

## Acknowledgments

We thank Bianca Splettstoesser for extensive technical support and Igor Paron, Christian Deiml, Johannes B. Müller, Antonio Piras, and Gabriele Sowa for technical assistance, Meera Phulphagar, Julia Schessner, and Georg Borner for their scientific input. Ashok Jayavelu provided scientific input and reagents, John Silke and Najoua Lalaoui (Walter and Eliza Hall Institute) provided Birinapant and compound A. Martin Spitaler and Markus Oster (Max Planck Institute of Biochemistry) helped with flow cytometry. Medini Steger provided helpful comments to the manuscript We thank all members of the Department of Proteomics and Signal Transduction at the Max Planck Institute of Biochemistry in Martinsried for their help and discussions, and especially

## Author contribution

M.C.T., M.M. conceived the project and wrote the manuscript. M.C.T. designed, performed all experiments, and analyzed all data (except ELISA). I.B. analyzed data and generated the website. C.A.S. performed the ELISA experiment and edited the manuscript. V.H. provided reagents and edited the manuscript.

## Methods

### Lead Contact and Materials Availability

Further information and requests for resources should be directed to and will be fulfilled by the Lead Contact, Matthias Mann. This study did not generate new unique materials and reagents.

### Experimental Model and Subject Details

U937 cells were purchased from ATCC and cultured in RPMI media with 10% FCS and Pen/Strep at 37 °C and 5% CO_2_. Mouse dermal fibroblasts (MDFs) were obtained from the epidermis and dermis of mouse tails through dispase II and collagenase digestion. Mouse dermal fibroblasts were then immortalized by infecting cells with the lentivirus SV40 Large T antigen. MDFs were cultured in DME media supplemented with 10% FCS and Pen/Strep at 37 °C and 5% CO_2_. Bone marrow-derived macrophages (BMDMs) were generated by flushing the bone marrow of hind bones with PBS. Cells were pelleted and were left for differentiation to macrophages in DME media with 20% L929 supernatant, non-essential amino acids, 10% FCS and Pen/Strep at 37 °C and 5% CO_2_ for 7 days. BMDMs were kept overnight in DME media with 10% FCS and Pen/Strep at 37 °C and 5% CO_2_ before stimulation.

### Method Details

#### Cell stimulation

U937 cells, BMDMs and MDFs were either left untreated or treated with TNF (100 ng/ml) alone or in combination of and IAP inhibitors/Smac mimetics (SM, Birinapant, 1.25 µM for U937 cells and Compound A, 2 µM for MDFs and BMDMs) to induce apoptosis or with TNF, SM and the pan-caspase inhibitor IDN-6556 (IDN-6556/Emricasan, 10 µM) to induce necroptosis. Cells were pretreated with inhibitors for one hour [RIPK1 inhibitor (necrostatin-1, 50 µM); TAK1 inhibitor (7-Oxozeaenol, 1 µM); IKK1/2 inhibitor (TPCA-1, 5 µM); p38 inhibitors (LY228820, 2 µM and SB203580, 10 µM); MEK inhibitor (PD0325901, 10 µM); JNK inhibitor (SP600125, 20 µM); TBK1/IKKε inhibitor (MRT67307, 2 µM); pan-CDK inhibitor (Dinaciclib: 6 nM for U937 cells, 12 nM for MDFs and 24 nM for BMDMs); CDK12/13 inhibitors (THZ531, 200 nM for U937 cells, and 400 nM for the proteome experiment described in Figure 5A, 800 nM for MDFs and BMDMs; SR4835, 60 nM for U937 cells and 160 nM for the qPCR experiment described in Figure 5 D, 160 nM for MDFs and 320 nM for BMDMs); CDK12 inhibitor (CDK12-IN3, 60 nM for U937 cells, 120 nM for MDFs and BMDMs), CDK9 inhibitors (Flavopiridol, 60 nM for U937 cells, MDFs and BMDMs; AZD4573, 6 nM for U937 and for MDFs and 24 nM for BMDMs). Stimulation details and inhibitor concentrations are also indicated in figure legends.

#### Phosphoenrichment protocol and proteome preparation for experiments and libraries

To enrich for phosphorylated peptides, we applied the Easy Phos protocol developed in our lab (Humphrey et al., 2015, Humphrey et al., 2018). In short, 10 × 10^6^ U937 cells or one full 15 cm dish of BMDMs were stimulated, washed three times with ice-cold TBS, lysed in 2% sodium deoxycholate (SDC) and 100 mM Tris-HCl [pH 8.5] and boiled immediately. After sonication, protein amounts were adjusted to 1 mg using the BCA protein assay kit. Samples were reduced with 10 mM tris(2-carboxy(ethyl)phosphine (TCEP), alkylated with 40 mM 2-chloroacetamide (CAA) and digested with Trypsin and LysC (1:100, enzyme/protein, w/w) overnight. For proteome measurements, 20 µg of the peptide was taken and desalted using SDB-RPS stage tips. 500 ng of desalted peptides were resolubilized in 5 µ*l* 2% ACN and 0.3% TFA and injected into the mass spectrometer. For phosphoenrichment, isopropanol (final conc. 50%), trifluoroacetic acid (TFA, final conc. 6%), and monopotassium phosphate (KH_2_PO_4_, final conc. 1 mM) were added to the rest of the digested lysate. Lysates were shaken, then spun down for 3 min at 2000 × g, and supernatants were incubated with TiO_2_ beads for 5 min at 40 °C (1:10, protein/beads, w/w). Beads were washed 5 times with isopropanol and 5% TFA, and phosphopeptides were eluted off the beads with 40% acetonitrile (ACN) and 15% of ammonium hydroxide (25% NH_4_OH) on C8 stage tips. After 20 min of SpeedVac at 45 °C, phosphopeptides were desalted on SDB-RPS stage tips and resolubilized in 5 µ*l* 2% ACN and 0.3% TFA and injected in the mass spectrometer.

To generate the library for phosphoproteome data independent acquisition (DIA) measurements, U937 cells were treated with TNF (100 ng/ml) in the presence of phosphatase inhibitors Sodium orthovanadate (1 mM) and Calyculin A (50 ng/ml) for 15 min. Cells were washed with ice-cold TBS and lysed in 2% SDC with 100 mM Tris-HCl [pH 8.5]. After boiling, sonication, reduction, alkylation, and overnight digestion (as described above), lysates were desalted on a Sepax Extraction column (Generik DBX), and 3 mg of desalted peptides were fractionated into 84 fractions on a C18 reversed-phase column (4.6 × 150 mm, 3.5 µm bead size) under basic conditions using a Shimadzu UFLX operating at 1 ml/minute. Peptides were separated on a linear gradient consisting of buffer A (2.5 mM Ammonium bicarbonate in MQ) and 2.5% to 44% buffer B (2.5 mM ABC in 80% ACN) for 64 min and 44% to 75% buffer B for 5 min before a rapid increase to 100%, which was kept for 5 min. Fractions were subsequently concatenated into 36 fractions and lyophilized. Phosphopeptides of these fractions were enriched using the phosphoenrichment protocol described above and run in data dependent acquisition (DDA) with the same gradient (70 min) the samples of the DIA experiments were run.

To generate the library for the proteome DIA measurements, U937 cells were lysed in 2% SDC with 100 mM Tris-HCl [pH 8.5], boiled, sonicated, reduced, alkylated, and digested overnight before desalting using SDB-RPS cartridges. 100 µ*g* of peptides were fractionated into 8 fractions by high pH reversed-phase chromatography. Fractions were concatenated automatically by shifting the collection tube every 120 s and subsequently dried in a SpeedVac. Peptides of these fractions were run in DDA mode with the same gradient (120 min) as the samples of the proteome DIA experiments.

#### Crude fractionation

After stimulation U937 cells (100 × 10^6^ cells) were washed with TBS and lysed in 4 ml lysis buffer (250 mM sucrose, 25 mM Tris-HCl [pH 7.5], 5 mM KCl, 3 mM MgCl_2_, 0.2 mM EDTA with protease- and phosphatase inhibitors (sodium pyrophosphate (12.5 mM = 50 × stock), β-glycerophosphate (2.2 g/10 ml = 1000 × stock), sodium orthovanadate (1.84 g/100 ml = 100 × stock)). Cells were broken up in the cell homogenizer (isobiotec) with 12 strokes. To 20% of lysate, SDS was added to a final concentration of 1% SDS and proteins were precipitated with acetone (final conc. 80%) for total phosphopeptide analysis. The rest of the lysate was spun down at 900 g for 10 min. The pellet represents the nucleus fraction and was resuspended in 1 % SDS before acetone precipitation. The supernatants were spun down for 40 min at 38.000 rpm using a Beckmann Coulter ultracentrifuge. The pellet, which contains the membrane fraction, was resuspended in 400 µ*l* of 1% SDS before the addition of acetone, while 1 ml of 2% SDS was added to the supernatant containing the cytosol fraction before protein precipitation with acetone. Proteins were precipitated overnight, and the next day washed with 80% acetone before resuspension in 4% SDC. The phosphoenrichment (1 mg/sample) and protein preparation (30 µ *g*/sample) of all fractions and conditions were performed as described above.

#### Western blotting

One million U937 cells were stimulated, washed in PBS, and lysed in buffer (1% IGEPAL, 10% Glycerol, 2 mM EDTA, 50 mM Tris pH 7.5, 150 mM NaCl) supplemented with protease inhibitors (Sigma-Aldrich, 4693159001). Lysates were kept on ice for 20 min and centrifuged at 16,100 × g for 15 min. After centrifugation 2 × SDS sample loading buffer (450 mM Tris-HCl, pH 8, 60% (v/v) glycerol, 12% (w/v) SDS, 0.02% (w/v) bromophenol blue, 600 mM DTT) was added to the supernatant before boiling of the lysate. Proteins were separated on 12% Novex Tris-glycine gels (Thermo Fisher Scientific, XP00120BOX) and transferred onto PVDF membranes (Merck Millipore, IPVH00010) or Nitrocellulose membranes (Amersham, 10600002). Membranes were blocked in 5% BSA in PBST, and antibodies were diluted in 2% BSA in PBST. Antibodies used for immunoblotting were as follows (diluted 1:1000): anti-human caspase-8 (MBL, M058-3), anti-cleaved human caspase-3 (Cell Signaling Technology CST, 9661), anti-human MLKL (Merck Millipore, MABC604), phospho anti-human RPB1 S2 (Millipore, 04-1571), anti-human RPB1 (CST, D8L4Y), phospho anti-human p65 (CST, 3033P), anti-human IκBα (CST, 9242), phospho anti-human p38 (CST, 9215), anti-human p38 (CST, 9212), anti-human CDK12 (CST, 11973), phospho anti-human CDK9 (CST, 2549), anti-human CDK9 (CST, 2316), anti-human FLIP (CST, 56343) and anti-human β-Actin (CST, 4967).

#### Elisa

1 x 10^4^ BMDMs were plated in 96 wells stimulated with CDK inhibitors for one hour before the addition of TNF (100 ng/ml). After 24 hours, the supernatants were spun down, and ELISA was performed according to the supplier’s protocol.

#### Cell death analysis

7 x 10^4^ U937 cells were plated in 96 wells, and BMDMs and MDFs were plated in 24 well plates and treated with TNF (100 ng/ml), IDN-6556 (10 µM), necrostatin-1 (Nec-1, 50 µM), Birinapant (SM, 1.25 µM for U937) and Compound A (SM, 2 µM for MDFs and BMDMs). Cell death was measured by propidium iodide incorporation using flow cytometry (FACS Attune NxT) and analyzed using Graphpad Prism.

#### qPCR

RNA was isolated from 2 x 10^6^ cells with the RNeasy Plus Mini kit (Qiagen) and reversely transcribed with SuperScript III (Invitrogen). cDNA was amplified with SYBR Green on a Biorad C1000 Thermal Cycler. Primers used were for MCP-1 (forward: CCTAGGAATCTGCCTGATAATCGA, reverse: TGGGATATACCATGCATACTGAGATG), and for GAPDH (forward: GTCTCCTCTGACTTCAACAGCG, reverse: ACCACCCTGTTGCTGTAGCCAA). Fold induction compared to untreated controls was calculated by the delta-delta CT method.

#### Chromatography and mass spectrometry

Samples were loaded onto 50-cm columns packed in-house with C18 1.9 μM ReproSil particles (Dr. Maisch GmbH), with an EASY-nLC 1000 system (Thermo Fisher Scientific) coupled to the MS (Q Exactive HFX, Thermo Fisher Scientific). A homemade column oven maintained the column temperature at 60°C. Peptides were introduced onto the column with buffer A (0.1% Formic acid), and phosphopeptides for data-independent acquisition were eluted with a 70 min gradient starting at 3% buffer B (80% ACN, 0.1% Formic acid) and followed by a stepwise increase to 19% in 40 min, 41% in 20 min, 90% in 5 min and 95% in 5 min, at a flow rate of 300 nL/min, while peptides for proteome analysis were eluted with a 120 min gradient starting at 5% buffer B (80% ACN, 0.1% Formic acid) followed by a stepwise increase to 30% in 95 min, 60% in 5 min, 95% in 2 x 5 min and 5% in 2 x 5 min at a flow rate of 300 nL/min. Phosphopeptides for data-dependent acquisition were eluted with a 140 min gradient starting at 5% buffer B (80% ACN, 0.1% Formic acid) followed by a stepwise increase to 20% in 85 min, 40% in 35 min, 65% in 10 min and 80% in 2 x 5 min at a flow rate of 300 nL/min.

A data-independent acquisition MS method was used for proteome and phosphoproteome analysis in which one full scan (300 to 1650 *m/z, R* = 60,000 at 200 *m/z*) at a target of 3 × 10^6^ ions was first performed, followed by 32 windows with a resolution of 30,000 where precursor ions were fragmented with higher-energy collisional dissociation (stepped collision energy 25%, 27.5%, 30%) and analyzed with an AGC target of 3 × 10^6^ ions and a maximum injection time at 54 ms in profile mode using positive polarity. Samples for libraries were run in data-dependent acquisition mode with the same gradients.

For the proteome/phosphoproteome libraries, samples were measured in data-dependent acquisition with a (TopN) MS method in which one full scan (300 to 1650/1600 *m/z, R* = 60,000 at 200 *m/z*) at a target of 3 × 10^6^ ions was first performed, followed by 15/10 data-dependent MS/MS scans with higher-energy collisional dissociation (target 10^5^ ions, maximum injection time at 28/60 ms, isolation window 1.4/1.6 *m/z*, normalized collision energy 27%, *R* = 15,000 at 200 *m/z*). Dynamic exclusion of 30 s was enabled.

For phosphoproteome measurements in data-dependent acquisition, a (TopN) MS method was used in which one full scan (300 to 1650 *m/z, R* = 60,000 at 200 *m/z*, maximum injection time 120 ms) at a target of 3 × 10^6^ ions was first performed, followed by 10 data-dependent MS/MS scans with higher-energy collisional dissociation (AGC target 10^5^ ions, maximum injection time at 120 ms, isolation window 1.6 *m/z*, normalized collision energy 27%, *R* = 15,000 at 200 *m/z*). Dynamic exclusion of 40 s and the Apex trigger from 4 to 7 s was enabled.

#### Quantification and Statistical Analysis

For the experiment measured in DDA mode, MS raw files were processed by the MaxQuant version 1.5.38 (Cox and Mann, 2008) and fragments lists were searched against the human UniProt FASTA database (21,039 entries, August 2015) (Cox et al., 2011) with cysteine carbamidomethylation as a fixed modification and N-terminal acetylation and methionine oxidations as variable modifications. For phosphoproteome analysis, we also added Serine/Threonine/Tyrosine phosphorylation as variable modification. We set the false discovery rate (FDR) to less than 1% at the peptide and protein levels and specified a minimum length of 7 amino acids for peptides. Enzyme specificity was set as C-terminal to Arginine and Lysine as expected using Trypsin and LysC as proteases and a maximum of two missed cleavages.

For experiments measured in DIA mode MS raw files were processed by the Spectronaut software version 13 (Biognosys, (Bruderer et al., 2015)). First, hybrid libraries were generated in Spectronaut Pulsar by combining the DDA runs of fractionated samples of either proteome or phosphoproteome with the DIA runs of the respective experiments. Human (21,039 entries, additional 74,013 entries, 2015) and mouse uniport FASTA databases (22,220 entries, 39,693 entries, 2015) as forward databases were used. To generate phosphoproteome libraries Serine/Threonine/Tyrosine phosphorylation was added as variable modification to the default settings which include cysteine carbamidomethylation as fixed modification and N-terminal acetylation and methionine oxidations as variable modifications. Maximum number of fragment ions per peptide was increased from 6 to 15. The false discovery rate (FDR) was set to less than 1% at the peptide and protein levels and a minimum length of 7 amino acids for peptides was specified. Enzyme specificity was set as C-terminal to Arginine and Lysine as expected using Trypsin and LysC as proteases and a maximum of two missed cleavages. To generate proteome libraries default settings were used. The experimental DIA runs were then analyzed against the hybrid library by using default settings for the analysis of the proteome and for the analysis of the phosphoproteome samples the localization cutoff was set to 0. BMDM raw files were analyzed via directDIA. For phosphoproteome analysis Serine/Threonine/Tyrosine phosphorylation was additionally set as variable modification and localization cutoff was set to 0.

All bioinformatics analyzes were done with the Perseus software (version 1.6.2.2) (Tyanova, 2016). For phosphosite analysis spectronaut normal report output tables were collapsed to phosphosites and the localization cutoff was set to 0.75 using the peptide collapse plug-in tool for Perseus (Bekker-Jensen et al., 2020). Summed intensities were log2 transformed. Samples that did not meet the measurement quality of the overall experiment were excluded. For the 8 min time point of the time course experiment we started with fewer replicates. Quantified proteins were filtered for at least 75% of valid values among three or four biological replicates in at least one condition. Missing values were imputed and significantly up- or down-regulated proteins were determined by multiple-sample test (one-way analysis of variance (ANOVA), FDR = 0.05) and Student’s t-test (two-sided), (FDR = 0.05). For **Figure 2G** we used two imputations to additionally obtain low abundant TNF-induced phosphosites (normal imputation: width 0.3, down shift 1.8, low imputation: width 0.15, down shift 3)

n represents replicates of the same cell line stimulated separately. Further statistical details of experiments can be found in the figure legends.

The 1D annotation enrichment analysis detects whether expression values of proteins belonging to an enrichment term (here we used: keywords, GOCC, GOMF, GOBP and KEGG name) show a systematic enrichment or de-enrichment compared to the distribution of all expression values (Cox and Mann, 2012).

Fisher’s exact tests were performed to detect the systematic enrichment or de-enrichment of annotations and pathways by analysing proteins whose levels or phosphorylation levels are significantly regulated upon different conditions (we used: keywords, GOCC, GOMF, GOBP and KEGG name). The Benjamini-Hochberg FDR represents the degree of significance and the enrichment factor the level of enrichment compared to the background.

Kinase motif enrichment analysis (**Figure 1I**) was performed by loading significantly (FDR < 0.05) up- and downregulated phosphosites on to the website: http://phosfate.com/profiler.html (Ochoa et al., 2016).

#### Dashboard for data visualization and manual inspection

For the TNF time course analysis and visualization, the median z-score and standard deviation for each phosphosite and time point were calculated per condition. The co-regulation analysis was performed by calculating all pairwise Pearson correlations between phosphopeptides across the sampled time points. The maximum q-value filter in the different analyses allows to filter for peptides or proteins that were significant in at least one ANOVA based comparison across conditions. A q-value of 0.01 means that maximally 1% of all results that are reported to be statistically significant are estimated to be false positives.

For sequence visualization and protein domain annotation, each phosphosite location was mapped to its respective protein sequence stored in the fasta file that was used for MS/MS data analysis (human fasta, downloaded 2015). The protein sequences for visualization were obtained using the ‘fasta’ functions from pyteomics (Goloborodko et al., 2013, Levitsky et al., 2019). Information about protein domains was retrieved from UniProt (https://www.uniprot.org/, accessed 22.06.2020 for human and 12.07.2020 for mouse), including following categories: ‘Chain’, ‘Domain’, ‘Alternative sequence’, ‘Propeptide’, ‘Signal peptide’ and ‘Transit peptide’.

Data preprocessing and visualization for the dashboard was performed using the python programming language. Following libraries were utilized for data processing: numpy, pandas, scipy, re and pyteomics (Goloborodko et al., 2013, Levitsky et al., 2019). Several libraries from the HoloViz family of tools were used for data visualization and creation of the dashboard, including panel, holoviews, param, bokeh, plotly and matplotlib.

---

**Figure S1:**
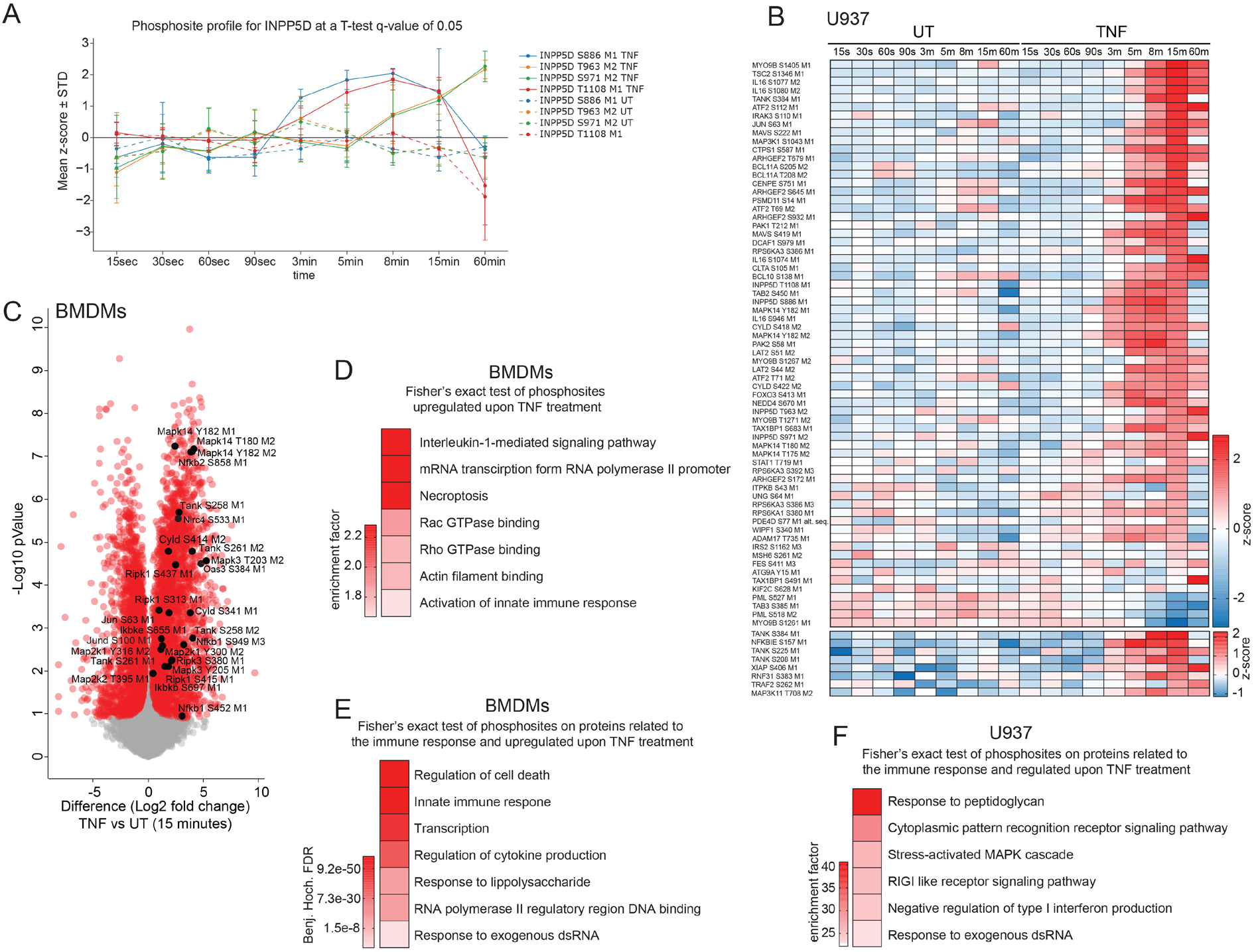
Members of other immune pathways are differentially phosphorylated upon TNF treatment. **A)** Mean of z-scored intensities of phosphosites detected on INPP5D (± SD). **B)** Heatmap of means of z-scored phosphosite intensities significantly changing in U937 cells upon TNF treatment compared to the untreated control filtered for the GOBP term ‘immune system process’ and various other GOBP enrichment terms, which include the term’ immune response’ (Student’s t-test FDR <0.05). **C**) Scatter plot of phosphosites regulated upon 15 min of TNF-treatment in BMDMs. Red dots represent significantly changing phosphosites (Student’s t-test FDR < 0.05). **D**) Fisher’s exact test of proteins with significantly increased phosphosites upon TNF treatment in BMDMs (p < 0.02). The following enrichment annotations are used: GOBP, GOCC, GOMF, keywords, and KEGG terms. **E**) Fisher’s exact test of proteins with upregulated phosphosites upon TNF stimulation filtered for the GOBP term’ immune system process’ and various other GOBP enrichment terms, which include the term ‘immune response’ (FDR < 0.02). The enrichment factor of all results was 26.2. **F**) Fisher’s exact test of proteins in **B**) compared to the whole dataset (FDR < 0.02).

**Figure S2:**
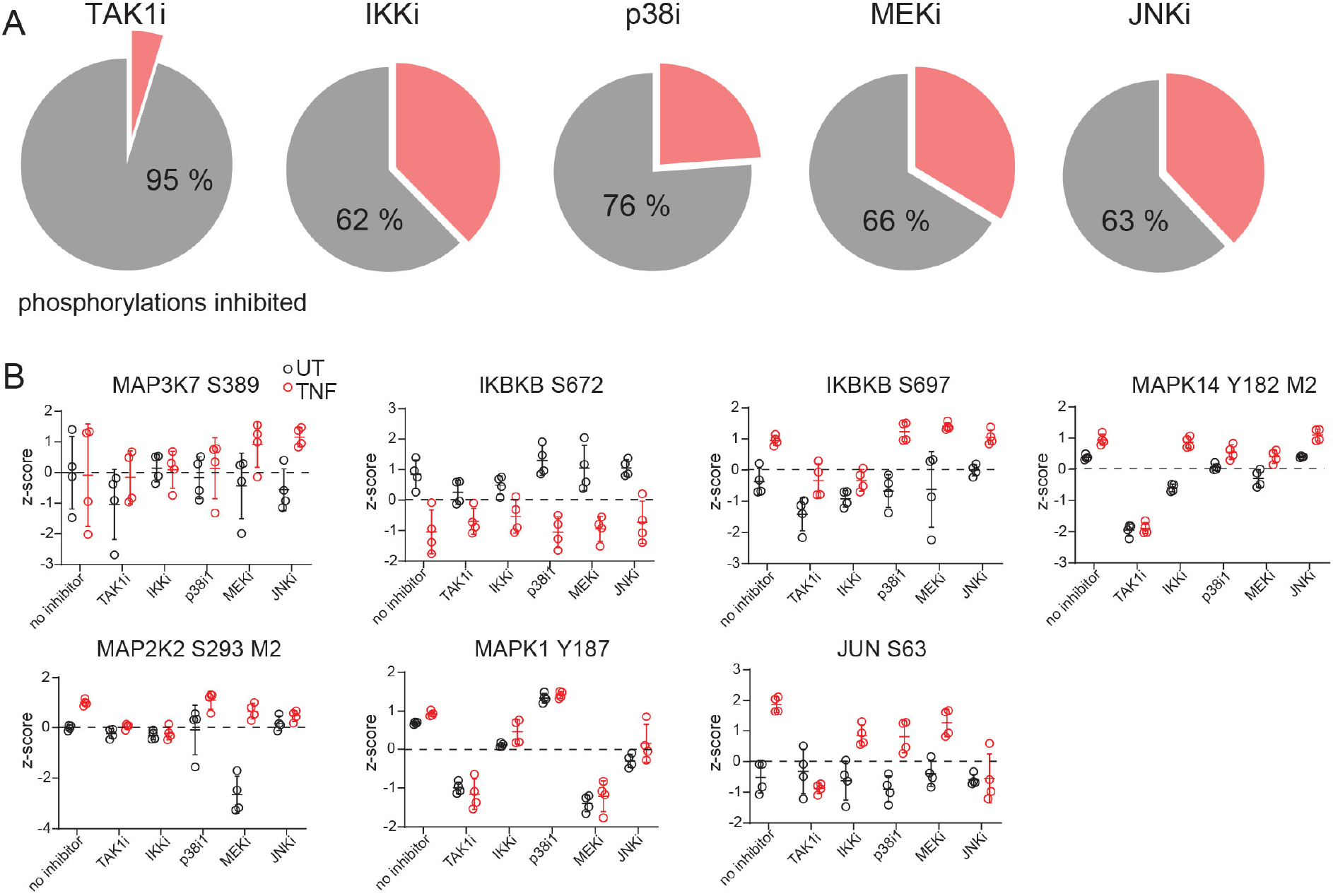
Kinase inhibitors have a major impact on the TNF-phosphoproteome. **A**) Pie charts of phosphosites significantly changing upon TNF treatment compared to the untreated control represent 100% (Student’s t-test FDR < 0.05). Inhibition of regulated phosphosites upon TNF treatment by different kinase inhibitors is shown in grey, while phosphosites still changing (Student’s t-test -Log10 p >2.0) upon the addition of kinase inhibitors are depicted in pink. **B**) Z-scored phosphosite intensities detected on known kinase substrates or kinases that are targeted by kinase inhibitors upon TNF treatment and in untreated conditions.

**Figure S3:**
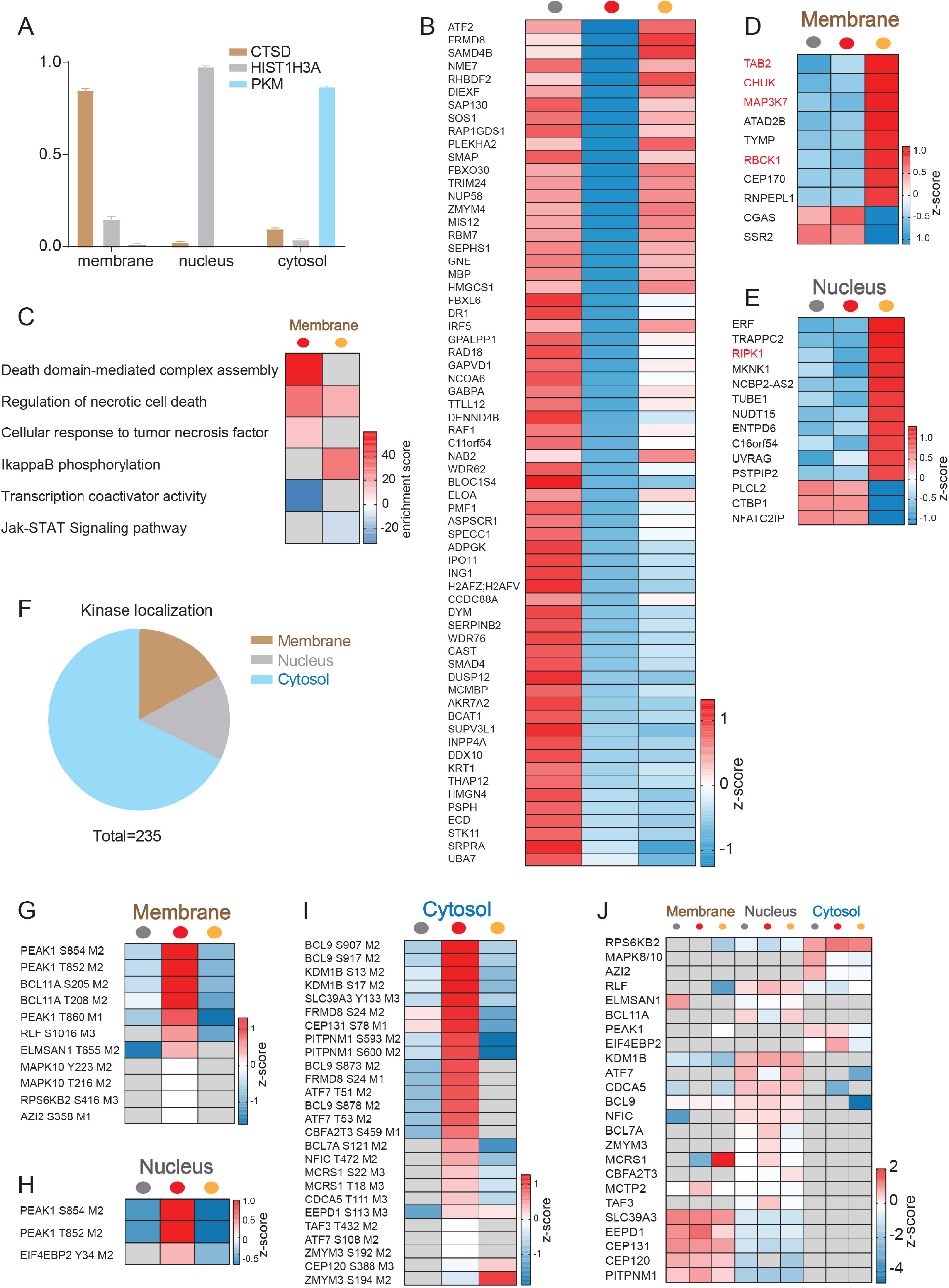
Protein translocation upon TNF-treatment. **A)** Normalized profiles of marker proteins of the membrane (CTSD), nuclear (HIST1H3A), and cytosolic (PKM) fractions throughout the three different fractions (± SD). **B**) Heatmap of means of z-scored protein intensities de-enriched in the membrane upon TNF treatment (Student’s t-test -Log10 p > 1.3, Log2 fold change < -0.5). **C**) Fisher’s exact test of proteins enriched and de-enriched (as shown in Figure 3D and Figure S3B) in the membrane fraction (p < 0.002). A negative enrichment score represents proteins downregulated upon treatment. **D**) Heatmap of z-score means of proteins changing in the membrane fraction upon TAK1 inhibition compared to untreated and TNF treated cells (Student’s t-test, -Log10 p > 1.3, Log2 fold change > 1.0 or < -1.0). **E**) Heatmap of means of z-scored protein intensities changing in the nucleus fraction upon TAK1 inhibition compared to untreated and TNF treated cells (Student’s t-test -Log10 p > 1.3, Log2 fold change > 1.0 or < -1.0). **F**) Pie chart representing the significant enrichment of kinases in the membrane, nucleus, and cytosol fraction (Student’s t-test FDR < 0.05). **G-I**) Heatmap of means of z-scored phosphosite intensities that are significantly changed in membrane **G**), nucleus **H**), and cytosol **I**) while respective proteins were not detected in the respective fractions (total valid values < 3). **J**) Heatmap of means of z-scored protein intensities carrying phosphosites that are significantly changed in membrane **G**), nucleus **H**), and cytosol **I**) while respective proteins were not detected in < 3 of 9 samples.

**Figure S4:**
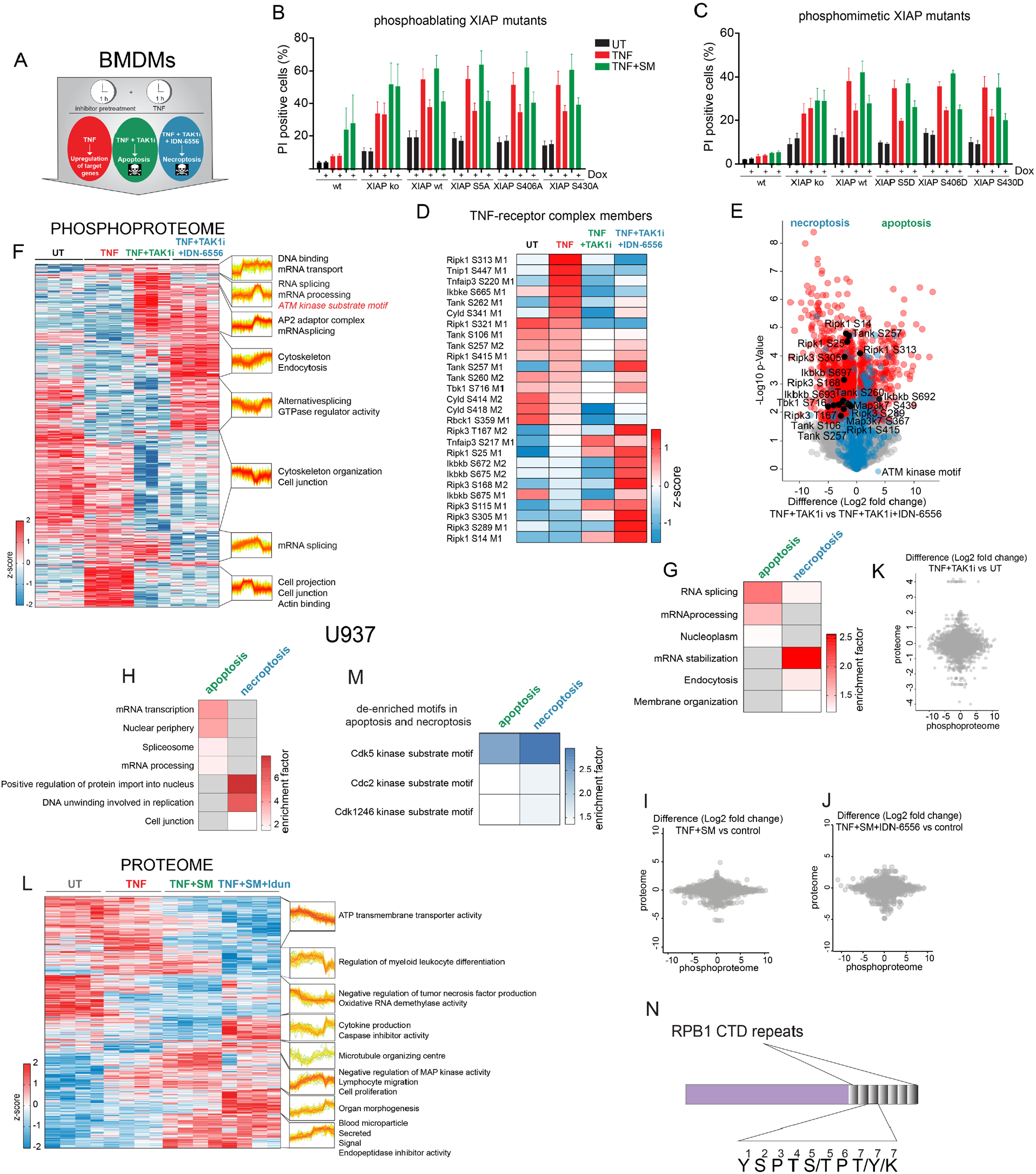
The phosphoproteome of BMDMs undergoing TNF-induced apoptosis and necroptosis. **A**) Experimental scheme of BMDMs treated with TNF (100 ng/ml, red), TNF and TAK1 inhibitor (TAK1i, 7-oxozeaenol, 1 μM, green) to induce apoptosis or with TNF, TAK1i and IDN-6556 (10 μM, blue) to induce necroptosis. **B-C**) Cell death analysis of propidium iodide stained U937 wt cells and XIAP deficient cells reconstituted with the indicated phosphoablative **B**) and mimetic **C**) XIAP mutants that were treated with TNF alone, TNF and Smac-mimetic (SM) or left untreated. XIAP expression was induced by the addition of doxycycline (1 μg/ml) (± SD, n ≥ 3). **D**) Heatmap of means of z-scored phosphosite intensities ANOVA significantly changing and localized on proteins associated with the TNF signaling complex (FDR < 0.05). **E**) Scatter plot of phosphosites regulated upon three hours of TNF-induced apoptosis compared to TNF-induced necroptosis in BMDMs. Red phosphosites are significantly changed (Student’s t-test FDR < 0.05). Blue phosphosites are part of an ATM kinase motif. **F**) Heatmap of z-scored phosphosite intensities ANOVA significantly changing upon TNF, TNF-induced apoptosis, or necroptosis treatment of BMDMs for three hours (FDR < 0.05). Fisher’s exact test of clusters of regulated phosphosites (FDR < 0.01). The profiles are color coded according to their distance from the respective cluster center (red is close to center, green is further away from center). **G**) Fisher’s exact test of phosphosites significantly increased (FDR < 0.05) upon apoptosis and necroptosis in BMDMs (p < 0.02). **H**) Fisher’s exact test of phosphosites of apoptotic and necroptotic U937 cells (p < 0.001). The red scale in G) and H) represents enrichment while grey shows no enrichment. **I-K**) Scatter plots of Student t-test difference comparisons of proteomes and respective phosphoproteomes of apoptotic and necroptotic U937 cells and apoptotic BMDMs. **L**) Heatmap of z-scored protein intensities ANOVA significantly changing upon TNF, TNF-induced apoptosis, or necroptosis treatment for three hours in U937 cells (FDR < 0.02). Fisher’s exact test of clusters of regulated proteins (p < 0.01). The profiles are color coded according to their distance from the respective cluster center (red is close to center, green is further away from center). **M**) Fisher’s exact test of downregulated phosphosite motifs of apoptotic and necroptotic U937 cells (FDR < 0.02). **N**) Scheme of RPB1 and example of CTD sequence repeats.

**Figure S5:**
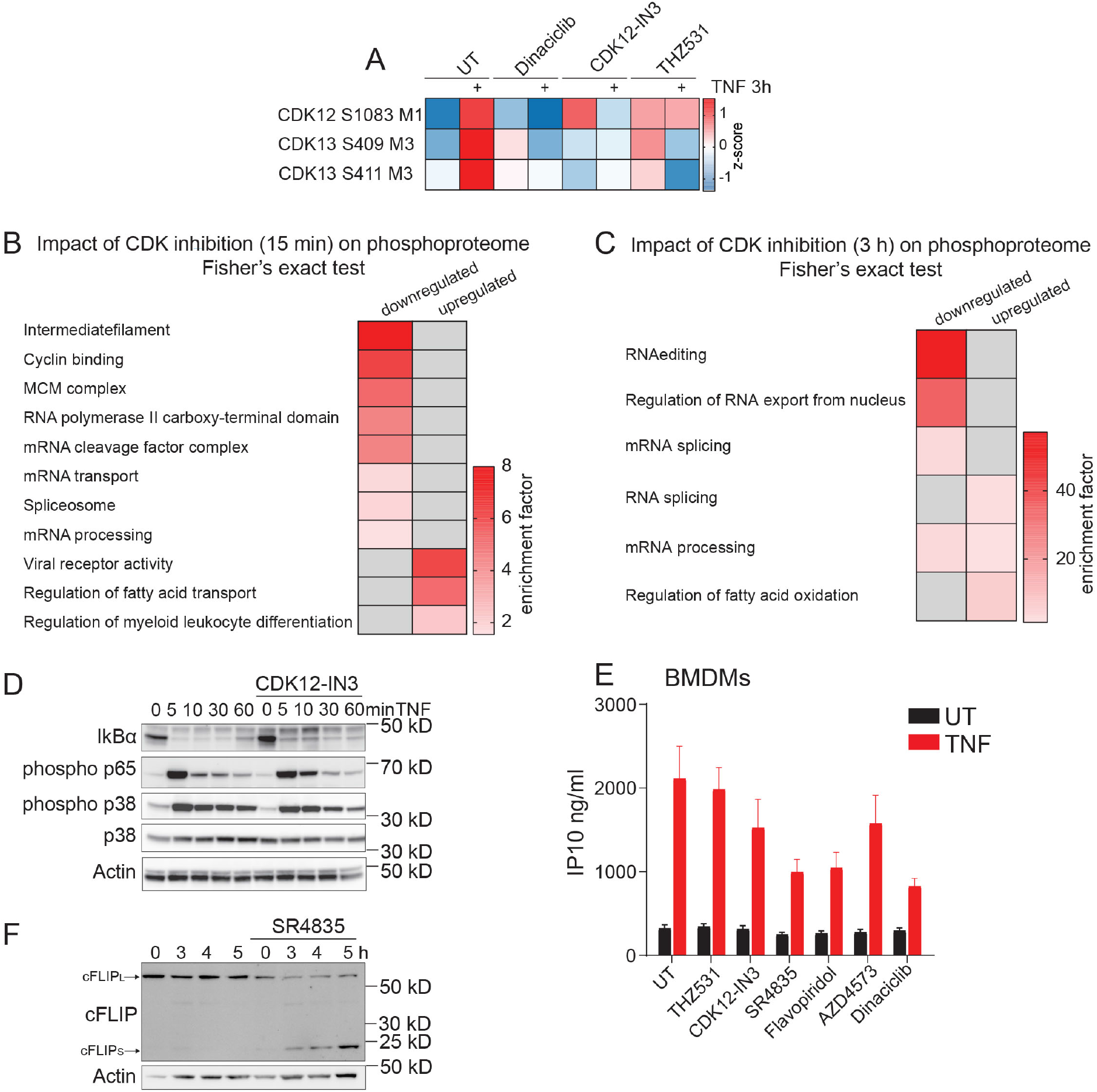
Inhibition of transcriptional CDKs impacts transcription and the release of TNF-induced cytokines. **A**) Heatmap of means of z-scored phosphosite intensities on CDK12 and CDK13 significantly changing upon 3 h of TNF stimulation (FDR < 0.05). **B**) Fisher’s exact test of down- and upregulated phosphosites upon CDK inhibition by Dinaciclib, CDK12-I, N3 and THZ531 for one hour and 15 min (p < 0.02). **C**) Fisher’s exact test of down- and upregulated phosphosites upon CDK inhibition by all three inhibitors tested: Dinaciclib, CDK12-I, N3, and THZ531 for four hours (p < 0.02 for downregulated, p < 0.002 for upregulated). **D**) Immunoblots of U937 cells treated with TNF alone and in combination with the CDK12 inhibitor CDK12-IN3 stained for members of the NFκB pathway, including IκBα, phosphorylated p65 and phosphorylated and total p38. **E**) ELISA of IP10 in the supernatant of BMDMs treated with or without TNF for 24 h, with and without CDK inhibitors (± SEM, n = 6). **F**) Immunoblot of U937 cells treated with TNF alone and in combination with CDK12/13 inhibitor SR4835 was stained for cFLIP and β-Actin.

**Figure S6:**
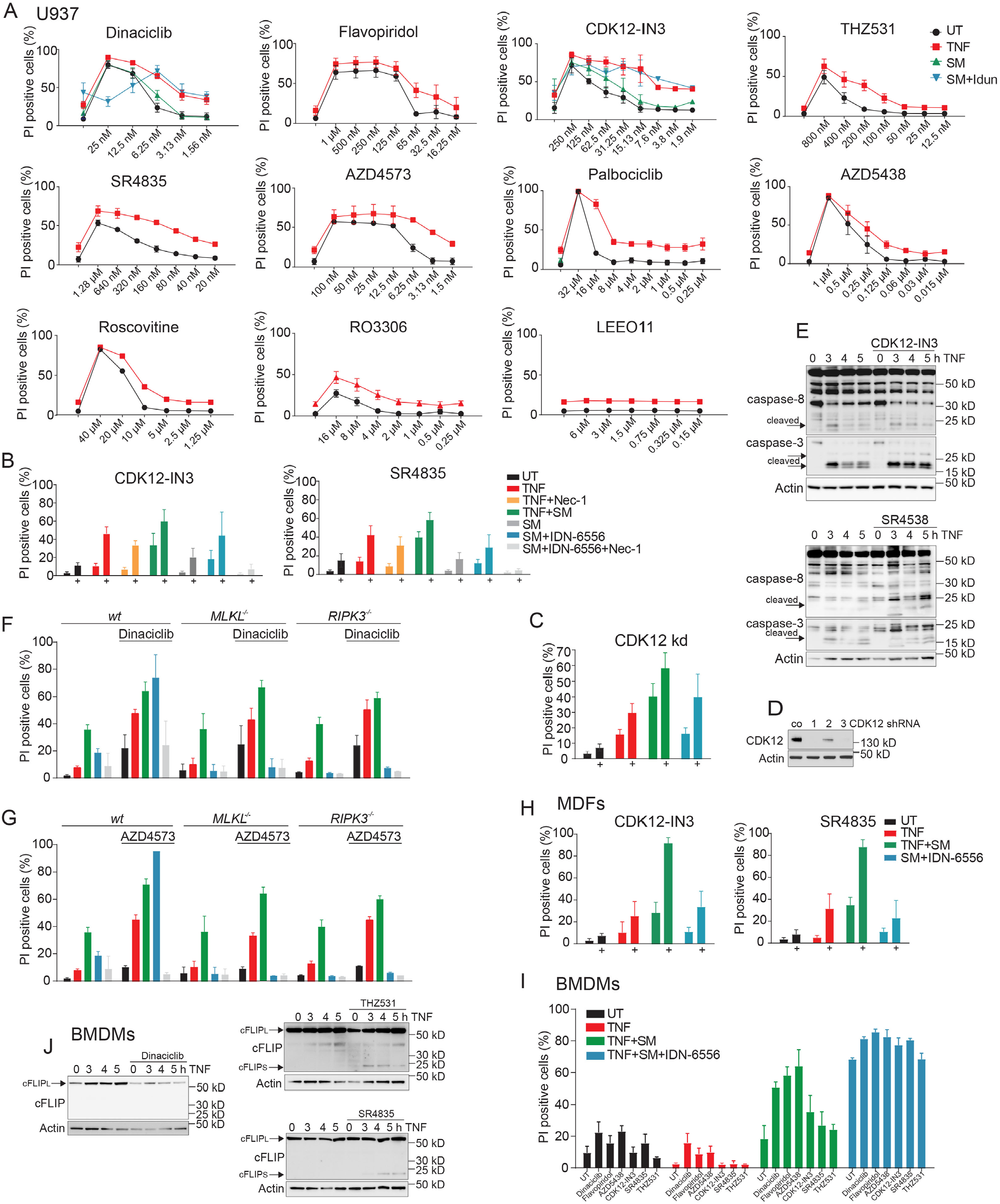
Various CDK inhibitors strongly enhance TNF-induced apoptosis and necroptosis. **A**) Cell death analysis of propidium iodide stained U937 cells treated as indicated over a concentration course of various CDK inhibitors for 24 h (± SD, n ≥ 2). **B**) Cell death analysis by flow cytometry of propidium iodide-stained U937 cells treated as indicated for 24 h (± SD, n ≥ 2). **C**) Cell death analysis by flow cytometry of propidium iodide stained U937 wt or CDK12 knocked down cells stimulated with TNF, TNF, and SM or TNF, SM, and IDN-6556 for 24 h (± SD, n ≥ 2). **D**) Immunoblot of U937 wt and CDK12 knockdown cells stained for CDK12 and β-Actin. **E**) Immunoblot of U937 cells stimulated with TNF alone for 3-5 h or in combination with CDK12-IN3 or SR4835 inhibitors and stained for caspase-3 and 8 and β-Actin. **F-G**) Cell death analysis of propidium iodide stained U937 wt cells or deficient for MLKL or RIPK3 treated as indicated for 24 h (± SD, n ≥ 2). **H**) Cell death analysis of propidium iodide stained MDFs treated as indicated for 24 h (± SD, n ≥ 2). **I**) Cell death analysis of propidium iodide stained BMDMs treated as indicated for 7 h (± SD, n ≥ 2). **J**) Immunoblots of BMDMs treated with TNF alone and in combination with CDK inhibitors were stained for cFLIP and β-Actin.

**Table S1: Phosphoproteome analysis of TNF-treated BMDMs**. BMDMs were treated with TNF for 15 min. This table reports phosphosites, the PTM localization probability within the peptide sequence, their fold change upon TNF stimulation and the significance (q-value).

## References

Bekker-Jensen, D. B., Bernhardt, O. M., Hogrebe, A., Martinez-Val, A., Verbeke, L., Gandhi, T., Kelstrup, C. D., Reiter, L. & Olsen, J. V. 2020. Rapid and site-specific deep phosphoproteome profiling by data-independent acquisition without the need for spectral libraries. Nat Commun, 11, 787.

Bradley, J. R. 2008. TNF-mediated inflammatory disease. J Pathol, 214, 149–60.

Brenner, D. W. & Shenderova, O. A. 2015. Theory and modelling of diamond fracture from an atomic perspective. Philos Trans A Math Phys Eng Sci, 373.

Bruderer, R., Bernhardt, O. M., Gandhi, T., Miladinovic, S. M., Cheng, L. Y., Messner, S., Ehrenberger, T., Zanotelli, V., Butscheid, Y., Escher, C., Vitek, O., Rinner, O. & Reiter, L. 2015. Extending the limits of quantitative proteome profiling with data-independent acquisition and application to acetaminophen-treated three-dimensional liver microtissues. Mol Cell Proteomics, 14, 1400–10.

Brumatti, G., Ma, C., Lalaoui, N., Nguyen, N. Y., Navarro, M., Tanzer, M. C., Richmond, J., Ghisi, M., Salmon, J. M., Silke, N., Pomilio, G., Glaser, S. P., De Valle, E., Gugasyan, R., Gurthridge, M. A., Condon, S. M., Johnstone, R. W., Lock, R., Salvesen, G., Wei, A., Vaux, D. L., Ekert, P. G. & Silke, J. 2016. The caspase-8 inhibitor emricasan combines with the SMAC mimetic birinapant to induce necroptosis and treat acute myeloid leukemia. Sci Transl Med, 8, 339ra69.

Budd, R. C., Yeh, W. C. & Tschopp, J. 2006. cFLIP regulation of lymphocyte activation and development. Nat Rev Immunol, 6, 196–204.

Cantin, G. T., Venable, J. D., Cociorva, D. & Yates, J. R., 3RD 2006. Quantitative phosphoproteomic analysis of the tumor necrosis factor pathway. J Proteome Res, 5, 127–34.

Cidado, J., Boiko, S., Proia, T., Ferguson, D., Criscione, S. W., San Martin, M., Pop-Damkov, P., Su, N., Roamio Franklin, V. N., Sekhar Reddy Chilamakuri, C., D’Santos, C. S., Shao, W., Saeh, J. C., Koch, R., Weinstock, D. M., Zinda, M., Fawell, S. E. & Drew, L. 2020. AZD4573 Is a Highly Selective CDK9 Inhibitor That Suppresses MCL-1 and Induces Apoptosis in Hematologic Cancer Cells. Clin Cancer Res, 26, 922–934.

Cox, J. & Mann, M. 2008. MaxQuant enables high peptide identification rates, individualized p.p.b.-range mass accuracies and proteome-wide protein quantification. Nat Biotechnol, 26, 1367–72.

Cox, J. & Mann, M. 2012. 1D and 2D annotation enrichment: a statistical method integrating quantitative proteomics with complementary high-throughput data. BMC Bioinformatics, 13 Suppl 16, S12.

Cox, J., Neuhauser, N., Michalski, A., Scheltema, R. A., Olsen, J. V. & Mann, M. 2011. Andromeda: a peptide search engine integrated into the MaxQuant environment. J Proteome Res, 10, 1794–805.

Croft, M., Benedict, C. A. & Ware, C. F. 2013. Clinical targeting of the TNF and TNFR superfamilies. Nat Rev Drug Discov, 12, 147–68.

Dondelinger, Y., Jouan-Lanhouet, S., Divert, T., Theatre, E., Bertin, J., Gough, P. J., Giansanti, P., Heck, A. J., Dejardin, E., Vandenabeele, P. & Bertrand, M. J. 2015. NF-kappaB-Independent Role of IKKalpha/IKKbeta in Preventing RIPK1 Kinase-Dependent Apoptotic and Necroptotic Cell Death during TNF Signaling. Mol Cell, 60, 63–76.

Dubbury, S. J., Boutz, P. L. & Sharp, P. A. 2018. CDK12 regulates DNA repair genes by suppressing intronic polyadenylation. Nature, 564, 141–145.

Feltham, R., Jamal, K., Tenev, T., Liccardi, G., Jaco, I., Domingues, C. M., Morris, O., John, S. W., Annibaldi, A., Widya, M., Kearney, C. J., Clancy, D., Elliott, P. R., Glatter, T., Qiao, Q., Thompson, A. J., Nesvizhskii, A., Schmidt, A., Komander, D., Wu, H., Martin, S. & Meier, P. 2018. Mind Bomb Regulates Cell Death during TNF Signaling by Suppressing RIPK1’s Cytotoxic Potential. Cell Rep, 23, 470–484.

Galbraith, M. D., Bender, H. & Espinosa, J. M. 2019. Therapeutic targeting of transcriptional cyclin-dependent kinases. Transcription, 10, 118–136.

Garber, M. E., Mayall, T. P., Suess, E. M., Meisenhelder, J., Thompson, N. E. & Jones, K. A. 2000. CDK9 autophosphorylation regulates high-affinity binding of the human immunodeficiency virus type 1 tat-P-TEFb complex to TAR RNA. Mol Cell Biol, 20, 6958–69.

Goloborodko, A. A., Levitsky, L. I., Ivanov, M. V. & Gorshkov, M. V. 2013. Pyteomics--a Python framework for exploratory data analysis and rapid software prototyping in proteomics. J Am Soc Mass Spectrom, 24, 301–4.

Greenleaf, A. L. 2019. Human CDK12 and CDK13, multi-tasking CTD kinases for the new millenium. Transcription, 10, 91–110.

Hayden, M. S. & Ghosh, S. 2008. Shared principles in NF-kappaB signaling. Cell, 132, 344–62.

Heger, K., Wickliffe, K. E., Ndoja, A., Zhang, J., Murthy, A., Dugger, D. L., Maltzman, A., De Sousa, E. M. F., Hung, J., Zeng, Y., Verschueren, E., Kirkpatrick, D. S., Vucic, D., Lee, W. P., Roose-Girma, M., Newman, R. J., Warming, S., Hsiao, Y. C., Komuves, L. G., Webster, J. D., Newton, K. & Dixit, V. M. 2018. OTULIN limits cell death and inflammation by deubiquitinating LUBAC. Nature, 559, 120–124.

Henry, K. L., Kellner, D., Bajrami, B., Anderson, J. E., Beyna, M., Bhisetti, G., Cameron, T., Capacci, A. G., Bertolotti-Ciarlet, A., Feng, J., Gao, B., Hopkins, B., Jenkins, T., Li, K., May-Dracka, T., Murugan, P., Wei, R., Zeng, W., Allaire, N., Buckler, A., Loh, C., Juhasz, P., Lucas, B., Ennis, K. A., Vollman, E., Cahir-Mcfarland, E., Hett, E. C. & Ols, M. L. 2018. CDK12-mediated transcriptional regulation of noncanonical NF-kappaB components is essential for signaling. Sci Signal, 11.

Hildebrand, J. M., Tanzer, M. C., Lucet, I. S., Young, S. N., Spall, S. K., Sharma, P., Pierotti, C., Garnier, J. M., Dobson, R. C., Webb, A. I., Tripaydonis, A., Babon, J. J., Mulcair, M. D., Scanlon, M. J., Alexander, W. S., Wilks, A. F., Czabotar, P. E., Lessene, G., Murphy, J. M. & Silke, J. 2014. Activation of the pseudokinase MLKL unleashes the four-helix bundle domain to induce membrane localization and necroptotic cell death. Proc Natl Acad Sci U S A, 111, 15072–7.

Hsu, H., Huang, J., Shu, H. B., Baichwal, V. & Goeddel, D. V. 1996. TNF-dependent recruitment of the protein kinase RIP to the TNF receptor-1 signaling complex. Immunity, 4, 387–96.

Humphrey, S. J., Azimifar, S. B. & Mann, M. 2015. High-throughput phosphoproteomics reveals in vivo insulin signaling dynamics. Nat Biotechnol, 33, 990–5.

Humphrey, S. J., Karayel, O., James, D. E. & Mann, M. 2018. High-throughput and high-sensitivity phosphoproteomics with the EasyPhos platform. Nat Protoc, 13, 1897–1916.

Hutti, J. E., Shen, R. R., Abbott, D. W., Zhou, A. Y., Sprott, K. M., Asara, J. M., Hahn, W. C. & Cantley, L. C. 2009. Phosphorylation of the tumor suppressor CYLD by the breast cancer oncogene IKKepsilon promotes cell transformation. Mol Cell, 34, 461–72.

Jaco, I., Annibaldi, A., Lalaoui, N., Wilson, R., Tenev, T., Laurien, L., Kim, C., Jamal, K., Wicky John, S., Liccardi, G., Chau, D., Murphy, J. M., Brumatti, G., Feltham, R., Pasparakis, M., Silke, J. & Meier, P. 2017. MK2 Phosphorylates RIPK1 to Prevent TNF-Induced Cell Death. Mol Cell, 66, 698–710 e5.

Johannes, J. W., Denz, C. R., Su, N., Wu, A., Impastato, A. C., Mlynarski, S., Varnes, J. G., Prince, D. B., Cidado, J., Gao, N., Haddrick, M., Jones, N. H., Li, S., Li, X., Liu, Y., Nguyen, T. B., O’Connell, N., Rivers, E., Robbins, D. W., Tomlinson, R., Yao, T., Zhu, X., Ferguson, A. D., Lamb, M. L., Manchester, J. I. & Guichard, S. 2018. Structure-Based Design of Selective Noncovalent CDK12 Inhibitors. ChemMedChem, 13, 231–235.

Kaczmarek, A., Vandenabeele, P. & Krysko, D. V. 2013. Necroptosis: the release of damage-associated molecular patterns and its physiological relevance. Immunity, 38, 209–23.

Krajewska, M., Dries, R., Grassetti, A. V., Dust, S., Gao, Y., Huang, H., Sharma, B., Day, D. S., Kwiatkowski, N., Pomaville, M., Dodd, O., Chipumuro, E., Zhang, T., Greenleaf, A. L., Yuan, G. C., Gray, N. S., Young, R. A., Geyer, M., Gerber, S. A. & George, R. E. 2019. CDK12 loss in cancer cells affects DNA damage response genes through premature cleavage and polyadenylation. Nat Commun, 10, 1757.

Krishnan, R. K., Nolte, H., Sun, T., Kaur, H., Sreenivasan, K., Looso, M., Offermanns, S., Kruger, M. & Swiercz, J. M. 2015. Quantitative analysis of the TNF-alpha-induced phosphoproteome reveals AEG-1/MTDH/LYRIC as an IKKbeta substrate. Nat Commun, 6, 6658.

Krystof, V., Baumli, S. & Furst, R. 2012. Perspective of cyclin-dependent kinase 9 (CDK9) as a drug target. Curr Pharm Des, 18, 2883–90.

Lafont, E., Draber, P., Rieser, E., Reichert, M., Kupka, S., De Miguel, D., Draberova, H., Von Massenhausen, A., Bhamra, A., Henderson, S., Wojdyla, K., Chalk, A., Surinova, S., Linkermann, A. & Walczak, H. 2018. TBK1 and IKKepsilon prevent TNF-induced cell death by RIPK1 phosphorylation. Nat Cell Biol, 20, 1389–1399.

Lee, D. F., Kuo, H. P., Chen, C. T., Hsu, J. M., Chou, C. K., Wei, Y., Sun, H. L., Li, L. Y., Ping, B., Huang, W. C., He, X., Hung, J. Y., Lai, C. C., Ding, Q., Su, J. L., Yang, J. Y., Sahin, A. A., Hortobagyi, G. N., Tsai, F. J., Tsai, C. H. & Hung, M. C. 2007. IKK beta suppression of TSC1 links inflammation and tumor angiogenesis via the mTOR pathway. Cell, 130, 440–55.

Lemke, J., Von Karstedt, S., Abd El Hay, M., Conti, A., Arce, F., Montinaro, A., Papenfuss, K., El-Bahrawy, M. A. & Walczak, H. 2014. Selective CDK9 inhibition overcomes TRAIL resistance by concomitant suppression of cFlip and Mcl-1. Cell Death Differ, 21, 491–502.

Levitsky, L. I., Klein, J. A., Ivanov, M. V. & Gorshkov, M. V. 2019. Pyteomics 4.0: Five Years of Development of a Python Proteomics Framework. J Proteome Res, 18, 709–714.

Ludwig, C., Gillet, L., Rosenberger, G., Amon, S., Collins, B. C. & Aebersold, R. 2018. Data-independent acquisition-based SWATH-MS for quantitative proteomics: a tutorial. Mol Syst Biol, 14, e8126.

Malumbres, M. 2014. Cyclin-dependent kinases. Genome Biol, 15, 122.

Mcilroy, D., Sakahira, H., Talanian, R. V. & Nagata, S. 1999. Involvement of caspase 3-activated DNase in internucleosomal DNA cleavage induced by diverse apoptotic stimuli. Oncogene, 18, 4401–8.

Messmann, R. A., Ullmann, C. D., Lahusen, T., Kalehua, A., Wasfy, J., Melillo, G., Ding, I., Headlee, D., Figg, W. D., Sausville, E. A. & Senderowicz, A. M. 2003. Flavopiridol-related proinflammatory syndrome is associated with induction of interleukin-6. Clin Cancer Res, 9, 562–70.

Micheau, O. & Tschopp, J. 2003. Induction of TNF receptor I-mediated apoptosis via two sequential signaling complexes. Cell, 114, 181–90.

Mihaly, S. R., Ninomiya-Tsuji, J. & Morioka, S. 2014. TAK1 control of cell death. Cell Death Differ, 21, 1667–76.

Mohideen, F., Paulo, J. A., Ordureau, A., Gygi, S. P. & Harper, J. W. 2017. Quantitative Phospho-proteomic Analysis of TNFalpha/NFkappaB Signaling Reveals a Role for RIPK1 Phosphorylation in Suppressing Necrotic Cell Death. Mol Cell Proteomics, 16, 1200–1216.

Murphy, J. M., Czabotar, P. E., Hildebrand, J. M., Lucet, I. S., Zhang, J. G., Alvarez-Diaz, S., Lewis, R., Lalaoui, N., Metcalf, D., Webb, A. I., Young, S. N., Varghese, L. N., Tannahill, G. M., Hatchell, E. C., Majewski, I. J., Okamoto, T., Dobson, R. C., Hilton, D. J., Babon, J. J., Nicola, N. A., Strasser, A., Silke, J. & Alexander, W. S. 2013. The pseudokinase MLKL mediates necroptosis via a molecular switch mechanism. Immunity, 39, 443–53.

Nakhaei, P., Sun, Q., Solis, M., Mesplede, T., Bonneil, E., Paz, S., Lin, R. & Hiscott, J. 2012. IkappaB kinase epsilon-dependent phosphorylation and degradation of X-linked inhibitor of apoptosis sensitizes cells to virus-induced apoptosis. J Virol, 86, 726–37.

O’Donnell, M. A., Perez-Jimenez, E., Oberst, A., Ng, A., Massoumi, R., Xavier, R., Green, D. R. & Ting, A. T. 2011. Caspase 8 inhibits programmed necrosis by processing CYLD. Nat Cell Biol, 13, 1437–42.

Oberst, A., Dillon, C. P., Weinlich, R., Mccormick, L. L., Fitzgerald, P., Pop, C., Hakem, R., Salvesen, G. S. & Green, D. R. 2011. Catalytic activity of the caspase-8-FLIP(L) complex inhibits RIPK3-dependent necrosis. Nature, 471, 363–7.

Ochoa, D., Jarnuczak, A. F., Vieitez, C., Gehre, M., Soucheray, M., Mateus, A., Kleefeldt, A. A., Hill, A., Garcia-Alonso, L., Stein, F., Krogan, N. J., Savitski, M. M., Swaney, D. L., Vizcaino, J. A., Noh, K. M. & Beltrao, P. 2020. The functional landscape of the human phosphoproteome. Nat Biotechnol, 38, 365–373.

Ochoa, D., Jonikas, M., Lawrence, R. T., El Debs, B., Selkrig, J., Typas, A., Villen, J., Santos, S. D. & Beltrao, P. 2016. An atlas of human kinase regulation. Mol Syst Biol, 12, 888.

Price, D. H. 2000. P-TEFb, a cyclin-dependent kinase controlling elongation by RNA polymerase II. Mol Cell Biol, 20, 2629–34.

Quereda, V., Bayle, S., Vena, F., Frydman, S. M., Monastyrskyi, A., Roush, W. R. & Duckett, D. R. 2019. Therapeutic Targeting of CDK12/CDK13 in Triple-Negative Breast Cancer. Cancer Cell, 36, 545–558 e7.

Salerno, D., Hasham, M. G., Marshall, R., Garriga, J., Tsygankov, A. Y. & Grana, X. 2007. Direct inhibition of CDK9 blocks HIV-1 replication without preventing T-cell activation in primary human peripheral blood lymphocytes. Gene, 405, 65–78.

Schmerwitz, U. K., Sass, G., Khandoga, A. G., Joore, J., Mayer, B. A., Berberich, N., Totzke, F., Krombach, F., Tiegs, G., Zahler, S., Vollmar, A. M. & Furst, R. 2011. Flavopiridol protects against inflammation by attenuating leukocyte-endothelial interaction via inhibition of cyclin-dependent kinase 9. Arterioscler Thromb Vasc Biol, 31, 280–8.

Silke, J. 2011. The regulation of TNF signalling: what a tangled web we weave. Curr Opin Immunol, 23, 620–6.

Sun, L., Wang, H., Wang, Z., He, S., Chen, S., Liao, D., Wang, L., Yan, J., Liu, W., Lei, X. & Wang, X. 2012. Mixed lineage kinase domain-like protein mediates necrosis signaling downstream of RIP3 kinase. Cell, 148, 213–27.

Taniguchi, C. M., Emanuelli, B. & Kahn, C. R. 2006. Critical nodes in signalling pathways: insights into insulin action. Nat Rev Mol Cell Biol, 7, 85–96.

Tanzer, M. C., Frauenstein, A., Stafford, C. A., Phulphagar, K., Mann, M. & Meissner, F. 2020. Quantitative and Dynamic Catalogs of Proteins Released during Apoptotic and Necroptotic Cell Death. Cell Rep, 30, 1260–1270 e5.

Ting, A. T., Pimentel-Muinos, F. X. & Seed, B. 1996. RIP mediates tumor necrosis factor receptor 1 activation of NF-kappaB but not Fas/APO-1-initiated apoptosis. EMBO J, 15, 6189–96.

Tyanova, S., Temu, T., Sinitcyn, P., Carlson, A., Hein, M., Geiger, T., Mann, M., Cox, J. 2016. The Perseus computational platform for comprehensive analysis of (prote)omics data. Nature Methods.

Wagner, S. A., Satpathy, S., Beli, P. & Choudhary, C. 2016. SPATA2 links CYLD to the TNF-alpha receptor signaling complex and modulates the receptor signaling outcomes. EMBO J, 35, 1868–84.

Welz, B., Bikker, R., Junemann, J., Christmann, M., Neumann, K., Weber, M., Hoffmeister, L., Preuss, K., Pich, A., Huber, R. & Brand, K. 2019. Proteome and Phosphoproteome Analysis in TNF Long Term-Exposed Primary Human Monocytes. Int J Mol Sci, 20.

Wertz, I. E. & Dixit, V. M. 2010. Regulation of death receptor signaling by the ubiquitin system. Cell Death Differ, 17, 14–24.

Wilson, N. S., Dixit, V. & Ashkenazi, A. 2009. Death receptor signal transducers: nodes of coordination in immune signaling networks. Nat Immunol, 10, 348–55.

Zhang, T., Kwiatkowski, N., Olson, C. M., Dixon-Clarke, S. E., Abraham, B. J., Greifenberg, A. K., Ficarro, S. B., Elkins, J. M., Liang, Y., Hannett, N. M., Manz, T., Hao, M., Bartkowiak, B., Greenleaf, A. L., Marto, J. A., Geyer, M., Bullock, A. N., Young, R. A. & Gray, N. S. 2016. Covalent targeting of remote cysteine residues to develop CDK12 and CDK13 inhibitors. Nat Chem Biol, 12, 876–84.

Zhong, C. Q., Li, Y., Yang, D., Zhang, N., Xu, X., Wu, Y., Chen, J. & Han, J. 2014. Quantitative phosphoproteomic analysis of RIP3-dependent protein phosphorylation in the course of TNF-induced necroptosis. Proteomics, 14, 713–24.

